# Transforming a head direction signal into a goal-oriented steering command

**DOI:** 10.1101/2022.11.10.516039

**Authors:** Elena A. Westeinde, Emily Kellogg, Paul M. Dawson, Jenny Lu, Lydia Hamburg, Benjamin Midler, Shaul Druckmann, Rachel I. Wilson

## Abstract

To navigate, we must continuously estimate the direction we are headed in, and we must use this information to guide our path toward our goal^1^. Direction estimation is accomplished by ring attractor networks in the head direction system^2,3^. However, we do not understand how the sense of direction is used to guide action. *Drosophila* connectome analyses^4,5^ recently revealed two cell types (PFL2 and PFL3) that connect the head direction system to the locomotor system. Here we show how both cell types combine an allocentric head direction signal with an internal goal signal to produce an egocentric motor drive. We recorded their activity as flies navigated in a virtual reality environment toward a goal stored in memory. Strikingly, PFL2 and PFL3 populations are both modulated by deviation from the goal direction, but with opposite signs. The amplitude of PFL2 activity is highest when the fly is oriented away from its goal; activating these cells destabilizes the current orientation and drives turning. By contrast, total PFL3 activity is highest around the goal; these cells generate directional turning to correct small deviations from the goal. Our data support a model where the goal is stored as a sinusoidal pattern whose phase represents direction, and whose amplitude represents salience. Variations in goal amplitude can explain transitions between goal-oriented navigation and exploration. Together, these results show how the sense of direction is used for feedback control of locomotion.

## Main

Accurate navigation requires us to fix a goal direction, and then maintain our orientation toward that goal in the face of perturbations. This is also a basic problem in mechanical engineering: how can we maintain the angle of some device at a particular goal value^6^? One solution to this problem is to use a resolver servomechanism to measure the discrepancy or error between the current angle and the goal angle. This produces a rotational velocity command that varies sinusoidally with error (Fig. 1a). Specifically, the mechanism drives leftward rotation when the device is positioned to the right of the goal, and *vice versa*.

**Figure 1:**
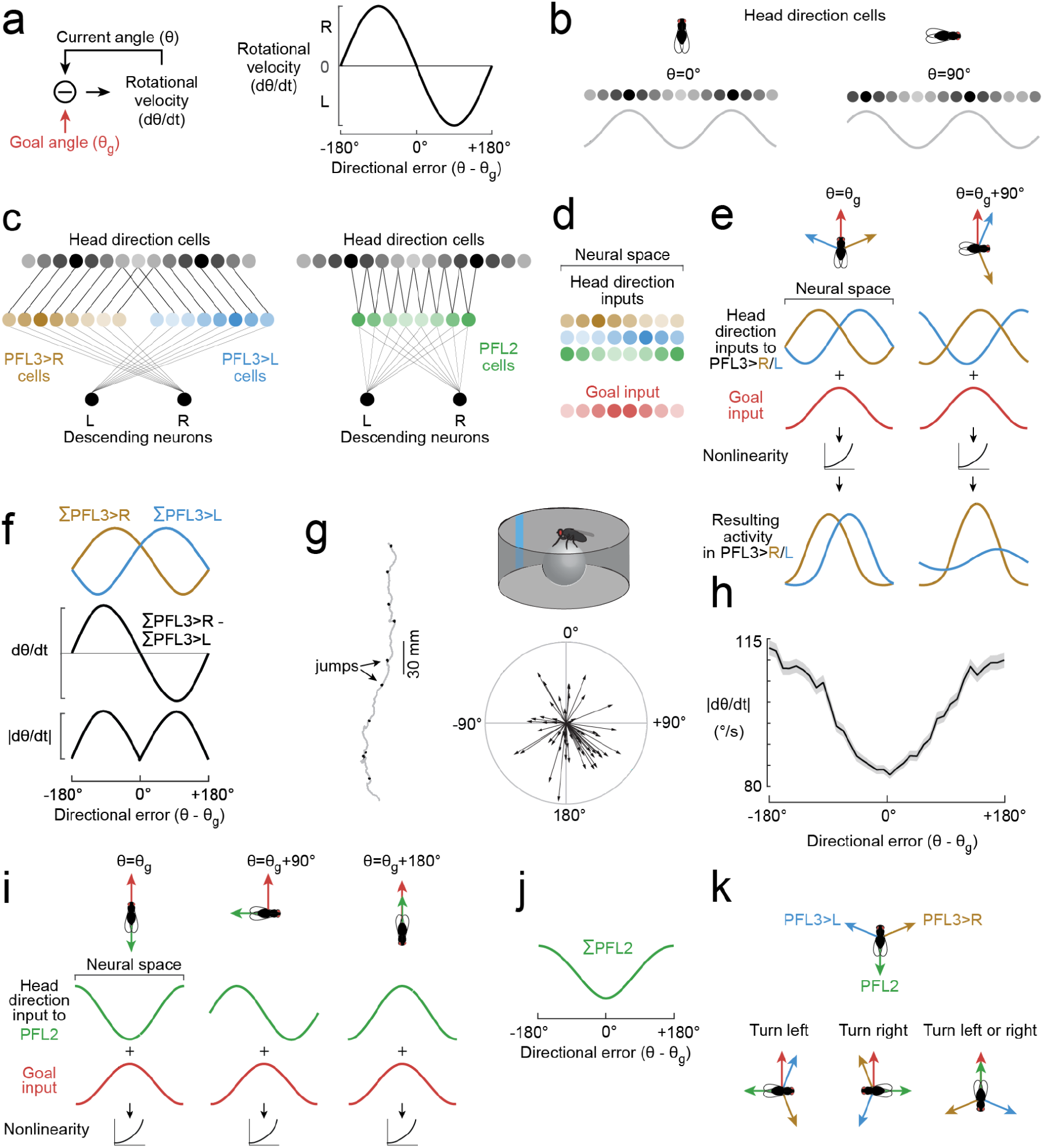
Comparing model predictions with behavior. **a**, A rotational servomechanism works to keep the angle of some device close to a goal value (θ_g_). The output of this device is a rotational velocity command, which can be a sinusoidal function of the system’s current error. Note that rotational velocity is close to zero around the goal (θ= θ_g_) and the anti-goal (θ= θ_g_+180°). **b**, In the *Drosophila* brain, head direction is represented in Δ7 cells as a sinusoid over two spatial cycles. Right turns push the phase left. **c**, PFL3>L, PFL3>R, and PFL2 populations extract shifted copies of this head direction representation. **d**, These three PFL populations are aligned in the fan-shaped body, where they share inputs from putative goal cells. The spatial phase of goal cell activity would shift when the fly’s internal goal changes. **e**, Model: PFL3 cells sum head direction input with goal input. This sum is passed through a nonlinearity. **f**, Model: total activity of each PFL3 population versus directional error (top). Right-left difference, taken as a predicted rotational velocity command (*ϱ*θ*ϱ*, middle). The absolute value of *ϱ*θ*ϱ*is rotational speed (bottom). **g**, Data: path of a fly in a virtual environment with a visual cue that rotates smoothly as the fly turns. Dots indicate abrupt jumps of the environment. Polar plot shows mean head direction in 10-min epochs, with radial length denoting the consistency of head direction over time (*ϱ*), n= 56 epochs from 56 flies; 0° is toward the cue. **h**, Data: mean rotational speed versus directional error. Bands are s.e.m. across flies (n=46 flies). **i**, Model: same as (e) but for PFL2. Here and in (e), neural activity is shown as a continuous function, but in reality it is discretized into 12 PFL3>R, 12 PFL3>L, and 12 PFL2 cells^5^. **j**, Model: total PFL2 activity versus directional error. **k** Model: PFL populations have head direction inputs that tile the space of compass directions. When their head direction input overlaps with the goal, they drive turning to reduce that overlap.

Sixty years ago, Mittelstaedt suggested that a similar process might be implemented in the brain’s navigation centers to control an organism’s heading, and thus its path through the environment^7^. Since then, several studies have proposed neural network implementations of this idea^4,5,8–14^. All these models exploit the fact that an angle or vector can be represented as a sinusoidal spatial pattern of activity across a neural population^15,16^. These sinusoids can then be combined to produce a directional control signal (Extended Data Fig. 1).

Data from locusts^17^, crickets^18^, zebrafish^19^, and *Drosophila*^*20*^ show that head direction is in fact encoded as a sinusoidal spatial pattern of activity (Fig. 1b). Interestingly, the *Drosophila* brain contains a cell type (PFL3) that is anatomically positioned to receive shifted copies of this head direction representation, while also making direct lateralized connections onto descending neurons involved in steering^4,5^ (Fig. 1c). This “copy-and-shift” architecture^21^ is reminiscent of the design of a resolver servomotor (Extended Data Fig. 2). PFL3 cells also receive anatomical input from the fan-shaped body, a brain region where goals might be stored (Fig. 1d). Notably, almost all the inputs to PFL3 cells are shared by another cell type, PFL2^5,22^. Individual PFL2 cells make bilateral connections onto descending neurons (Fig. 1c), implying that they do not guide steering. Their function is enigmatic, but they have been proposed to increase forward walking speed^5,11^.

In short, both PFL2 and PFL3 cells are anatomically positioned to integrate current spatial information with stored goal information for navigation control. These cells stand out because they form a link between an allocentric map of space and an egocentric system of motor control. However, there have been no functional studies of these cells, and recent models of this network have made conflicting predictions^4,5,10,11,13^.

### Comparing model predictions with behavior

To begin, we describe an updated computational model based on connectome data^4,5^. Our model shares some features with other recent models^4,5,10,11,13^, but some of our key predictions are different (see Methods: Network model). This is partly because we use automatic neurotransmitter assignments^23^ to constrain the signs of each connection in the network.

In this model, head direction is represented as a sinusoid^24^ whose phase tracks the fly’s head direction, relative to a flexible and arbitrary offset^20^. We divide PFL3 cells into two populations (PFL3>R and PFL3>L) that converge onto right or left descending neurons, respectively (Fig. 1c). Each population extracts a copy of the head direction representation, with phase ±67.5°-θ_0_, where θ_0_ is an arbitrary reference value. Meanwhile, PFL2 cells extract a head direction representation with phase 180°-θ_0_ (Fig. 1c).

These three PFL populations are aligned within the fan-shaped body, where they share inputs from orderly arrays of cells^5^ which could represent the goal angle, θ_g_ (Fig. 1d). We model the goal representation as a spatial sinusoid with phase (θ_g_-θ_0_). The goal might be stored as sinusoidal pattern of persistent activity in the network, or a sinusoidal pattern of synaptic weights. The firing rate of each model PFL cell is the sum of its head direction input and its goal input, passed through a nonlinearity (Fig. 1e).

It should be remembered that these sinusoids are representations of vectors (Extended Data Fig. 1). In essence, the two PFL3 populations extract shifted copies of the head direction vector, and the goal vector is added to each copy. The vector with the larger magnitude dictates the direction the fly should rotate to reach its goal.

This model predicts that PFL3>R should be most active when the fly is facing to the left of its goal, and *vice versa* for PFL3>L (Fig. 1f). PFL3 cells are excitatory (see Methods) and they directly excite descending neurons involved in steering whose right-left difference drives proportional increases in rotational velocity^4^; thus, the direct PFL3 contribution to rotational velocity should be ΣPFL3>R - ΣPFL3>L (Fig. 1f).

Note that, in this model, PFL3 cells cause the fly to behave like a resolver servomechanism: rotational velocity is low around the goal, and also opposite the goal (Fig. 1f). Engineers call this “false nulling”, because it can allow a servomechanism to settle at an angle opposite the goal. To look for this phenomenon in fly behavior, we placed flies in a virtual reality environment with a single prominent visual head direction cue; this environment rotated in closed loop with the fly’s rotational velocity on a spherical treadmill (Fig. 1g). The fly’s head was rigidly coupled to its body, so that heading and head direction are identical. In this type of environment, flies often follow straight paths toward a goal stored in memory, and this behavior requires an intact head direction system^25–27^ (Fig. 1i). During these epochs of straight walking, we could infer the fly’s goal direction from its behavioral orientation. Every few minutes, we jumped the virtual environment by 90° or 180°; this generally caused the fly to turn back toward its goal (Fig. 1g), implying that these jumps are perceived by the brain as head direction changes^4,25^. In agreement with our model predictions, we found that the fly’s rotational speed was generally minimal when it was oriented toward its goal. However, contrary to predictions, the fly’s rotational speed was high — not low — around its anti-goal, and 180° jumps evoked rotational speeds that were no lower than those evoked by 90° jumps (Extended Data Fig. 3a). Our PFL3 model cannot explain this result (Fig. 1f), suggesting an additional mechanism recruited around the anti-goal to promote rapid re-orientation. PFL2 cells are good candidates for this mechanism, because their amplitude should be highest when the fly is oriented toward its anti-goal (Fig. 1i,j). If PFL2 cells promote turning away from the anti-goal, this would solve the false-nulling problem, ensuring that the fly would not walk straight in the direction opposite its goal.

In summary, our model predicts that all these PFL populations have head direction inputs that tile the space of compass directions (Fig. 1k). When their head direction input overlaps with the goal input, each population drives a motor response to reduce that overlap. Together, these cells should keep the fly’s path generally aligned with its goal, even in the face of perturbations or obstacles that require a momentary detour.

### Dynamics around the anti-goal

To test our model predictions, we constructed split-Gal4 lines to target PFL2 and PFL3 cells. We were able to generate a highly selective PFL2 line, as well as a line targeting PFL2 and PFL3 together. We validated these lines by using genetic mosaic analysis to identify single-cell clones, and then comparing these clones to morphologies from the connectome (Extended Data Fig. 4a,c). We will focus initially on our results for PFL2 cells, as this line was the more specific one.

First, to directly activate PFL2 cells, we used a chemogenetic approach: we expressed ATP-gated ion channels (P2×2 receptors) in these cells, and we activated them specifically using iontophoresis of ATP into the protocerebral bridge, where their dendrites are located (Fig. 2a, Extended Data Fig. 5). We made a whole-cell recording from a PFL2 cell in every experiment to confirm that ATP produced precise and transient depolarization (Fig. 2a,b). At the same time, we monitored the fly’s behavior on a spherical treadmill, again a virtual reality environment with a visual cue. As predicted, we found that stimulating PFL2 cells generally produced turning, although the direction of the turn was often unpredictable (Fig. 2a,b). Moreover, if the fly was walking forward at the time of the stimulus, it consistently reversed direction and stepped backward (Fig. 2a,b). This response may be related to the fact that bidirectional excitation in some steering-related descending neurons is correlated with slower forward movement, or even backward locomotion^4^. Together, these results show that PFL2 cells drive an increase in rotational movement, accompanied by a decrease in forward velocity.

**Figure 2:**
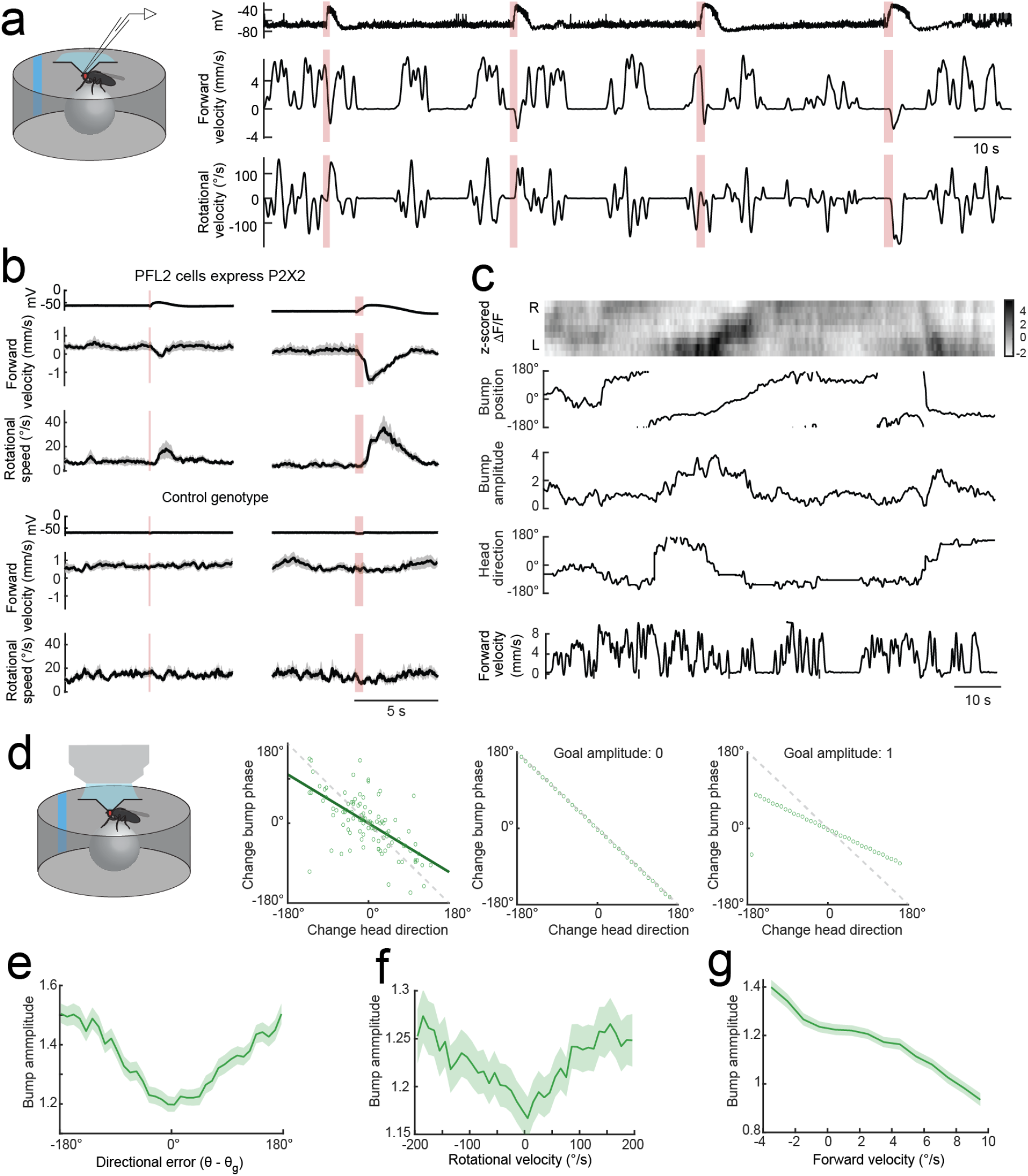
Dynamics around the anti-goal. **a**, Example experiment. ATP iontophoresis depolarizes PFL2 cells expressing P2×2 (top). This stimulus evokes an increase in rotational speed (middle) and a decrease in forward velocity (bottom). This fly turns right in response to the first two pulses, but left in response to the second two pulses. **b**, Summary data for flies where PFL2 cells expressed P2×2 and genetic controls (mean ± s.e.m. across flies, n=12 P2×2+ flies and 11 control flies). Results are shown for two ATP pulse durations (100 ms and 500 ms). In flies where PFL2 cells expressed P2×2, ATP also increases sideways speed (Extended Data Fig. 5). **c**, PFL2 activity (ΔF/F) across the horizontal axis of the fan-shaped body, over time. During this epoch, the fly is walking relatively straight. The fly’s mean head direction is taken as its angular goal (θ_g_). After the environment is jumped by 180°, the fly makes a compensatory turn to reorient toward θ_g_. We fit a sinusoid to ΔF/F at each time point to extract phase and amplitude. **d**, Change in PFL2 phase versus change in head direction, over successive imaging volumes. Head direction values are expressed relative to the fly’s goal. The phase of PFL2 activity moves right when the fly turns left. Each symbol denotes one time point (Pearson’s R = −0.63, p = 9×10^−13)^,. For comparison, model results are shown for several different values of the goal amplitude (*A*). Shown here are data for one example fly; Extended Data Fig. 8 shows more examples. **e**, Amplitude of PFL2 activity versus directional error (mean ± s.e.m. across flies, n=33 flies). The mean amplitude is highest around the anti-goal. **f**, Amplitude of PFL2 activity versus the fly’s rotational speed (mean ± s.e.m. across flies, n=33 flies). **g**, Amplitude of PFL2 activity versus the fly’s forward velocity (mean ± s.e.m. across flies, n=33 flies).

Next, we asked when PFL2 cells are active. We used PFL2-split-Gal4 to drive expression of GCaMP7b, and we imaged the activity of these cells with a 2-photon microscope. As before, the fly navigated in a virtual environment with a visual cue, and we inferred the fly’s goal direction from its behavioral orientation during epochs of voluntary straight-line walking. Every few minutes, we jumped the environment by 90° or 180°; thus forcing flies to sample a range of directions.

We saw that PFL2 activity (ΔF/F) generally formed a sinusoidal spatial pattern across the horizontal axis of the fan-shaped body (Fig. 2c). Therefore, we fit a sinusoidal function to these patterns, and we extracted the phase and amplitude of this activity, which we refer to as the “amplitude” of neural activity. We found that the phase of PFL2 activity generally moved leftward as the fly rotated to the right (Fig. 2d), as predicted.

Notably, we found that the amplitude of PFL2 activity was minimal when the fly was oriented toward its goal, and maximal around the anti-goal (Fig.2e). Moreover, the amplitude of PFL2 activity was predictive of the fly’s behavior: high amplitude predicted high rotational speed (Fig. 2f) and low forward velocity (Fig. 2g). Together, these data show that PFL2 cells are recruited when the fly is facing its anti-goal, driving a nondirectional increase in turning, accompanied by a decrease in forward velocity. Thus, these cells provide a solution to the “false nulling” problem that characterizes a classical servomechanism.

### Dynamics around the goal

Next, we imaged GCaMP7b expressed under the control of the mixed split-Gal4 line that targets both PFL2 and PFL3 cells (Extended Data Fig. 4). Here, rather than focusing on the fan-shaped body, we focused on the lateral accessory lobes, where PFL axons terminate, in order to separate PFL3>L from PFL3>R. PFL2 and PFL3 axon terminals are intermingled in the lateral accessory lobes, but we found that calcium signals in the mixed line were quite different from the signals we observed in PFL2 cells. In the PFL2-specific line, calcium signals in the lateral accessory lobes were generally maximal around the anti-goal (Fig. 3a), as we would expect from our imaging data in the fan-shaped body and protocerebral bridge (Fig. 2e). However, in the mixed line, we saw the opposite: calcium signals in the lateral accessory lobes were generally maximal around the goal (Fig. 3b). This is what we would expect from the PFL3 populations, and it implies that the signals in the mixed line are dominated by PFL3 cells. This could be due to stronger Gal4 expression in PFL3 versus PFL2, or intrinsic cell-specific features that cause axonal GCaMP7b signals to be dominated by PFL3. Thus, we can treat the lateral accessory lobe signals as a readout of the summed activity of each PFL3 population.

**Figure 3:**
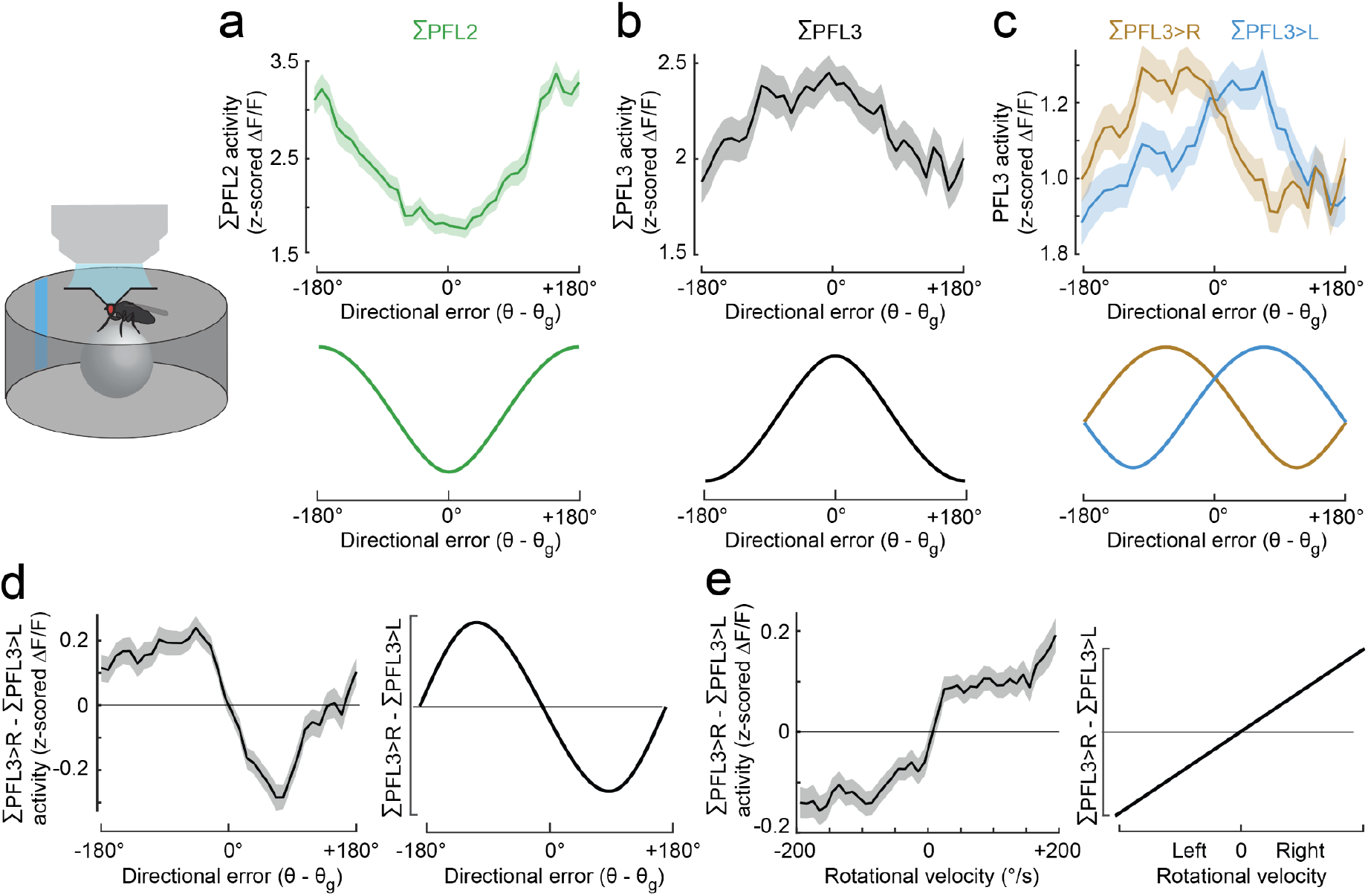
Dynamics around the goal. **a**, PFL2 activity (ΔF/F) in the lateral accessory lobe. Shown here is the summed activity of the right and left PFL2 axons, versus directional error (mean ± s.e.m. across flies, n=33 flies). The model prediction is shown for comparison. **b**, PFL2+3 activity (ΔF/F) summed across the right and left lateral accessory lobe, versus directional error (mean ± s.e.m. across flies, n=23 flies). Here we used a mixed split-Gal4 line that targets PFL2 and PFL3 cells together. Results are opposite for what we observe for PFL2 cells alone, and it matches the predictions of our PFL3 model (shown for comparison), implying that PFL3 dominates ΔF/F. **c**, ΣPFL3>R and ΣPFL3>L activity in the right and left lateral accessory lobe, versus directional error (mean ± s.e.m. across flies, n=23 flies). Signals are imaged from our mixed split-Gal4 line but are likely dominated by PFL3. Predicted responses of model PFL3 cells are shown for comparison. **d**, Right-left difference in PFL3 activity versus directional error (mean ± s.e.m. across flies, n=23 flies), and model prediction. **e**, Right-left difference in PFL3 activity versus the fly’s rotational velocity (mean ± s.e.m. across flies, n=23 flies), and model prediction.

In agreement with our model predictions, we found that PFL3>R cells are most active when the fly is just to the left of its goal, and *vice versa* for PFL3>L (Fig. 3c).The difference between these populations (ΣPFL3>R - ΣPFL3>L) is a roughly sinusoidal function of the fly’s orientation relative to its goal, supporting the predictions of our model (Fig. 3d). Moreover, we found that the right-left difference in PFL3 activity was predictive of the fly’s rotational velocity, again consistent with our model (Fig. 3g). Unlike PFL3 cells, PFL2 axonal projections were symmetrically active regardless of head direction, as we would predict based on their anatomy (Extended Data Fig. 6a).

In summary, our data argues that PFL3 cells drive precise directional steering maneuvers that correct small deviations from the fly’s intended path. PFL3 cells are typically recruited when the fly is near its goal direction. This stands in contrast to PFL2 cells, which are recruited primarily around the anti-goal.

### Generating behavioral variations

Next, we used connectome data to make behavioral predictions based on our network model. PFL3 cells make direct excitatory connections onto steering descending neurons (DNa02); they also connect to DNa03, which is one of the strongest excitatory inputs to DNa02^4,5,28,29^. Thus, PFL3 cells are positioned to influence steering via a direct pathway and an indirect pathway (Fig. 4a). PFL2 cells do not connect directly to DNa02, but they make a massive connection onto DNa03, meaning they are positioned to control signaling through the indirect pathway (Fig. 4a).

**Figure 4:**
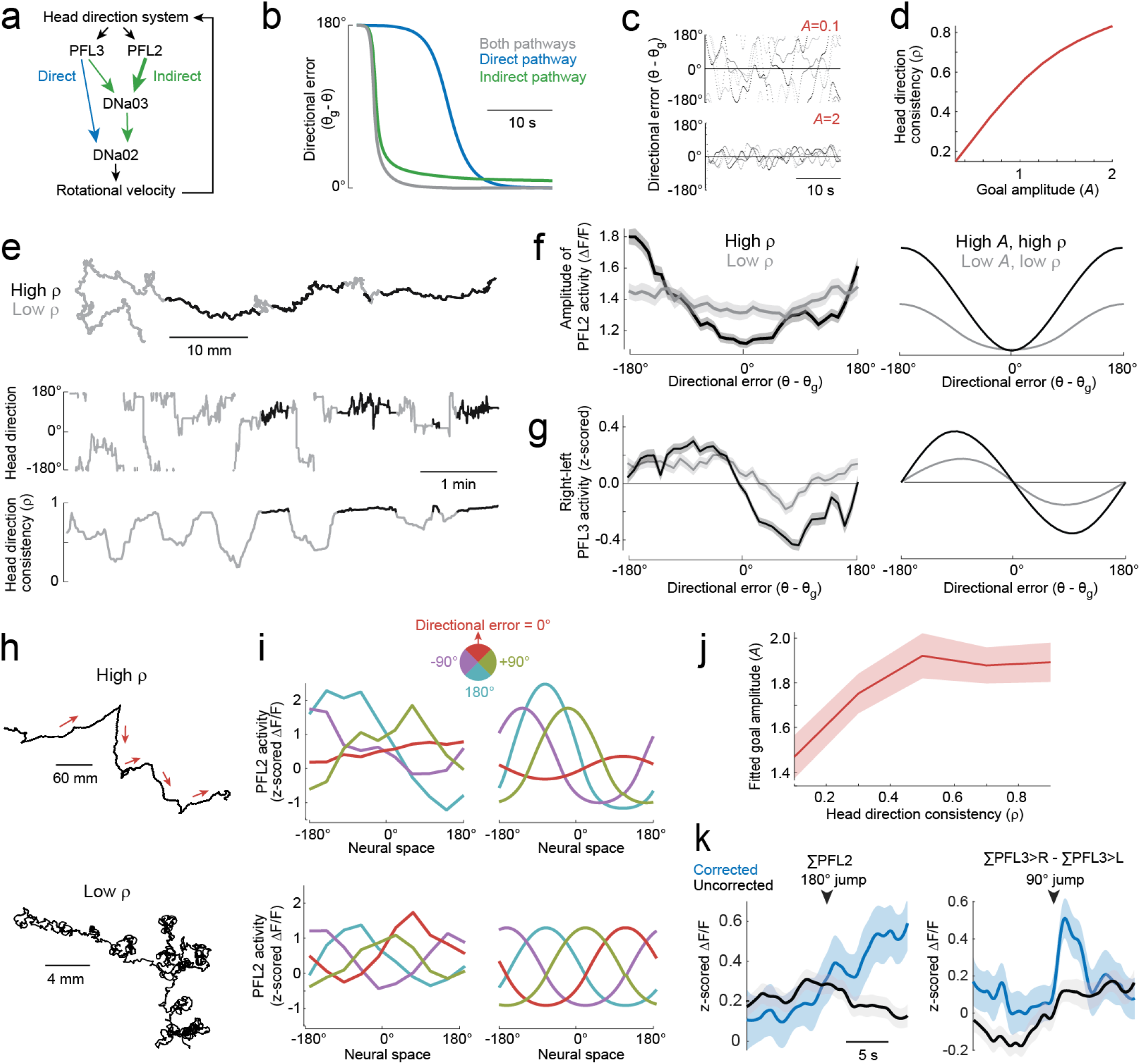
Generating behavioral modulations. **a**, Direct pathway and indirect pathways. We model the influence of PFL2 cells as a multiplicative scaling of the indirect pathway. **b**, Model: effect of each pathway alone and in combination. Directional error versus time after the fly is rotated 180° away from the goal. **c**, Model: head direction (relative to goal) over time, for 2 values of goal amplitude *A*. Independent noise was injected into the steering signal on each trial. **d**, Model: mean head direction consistency versus goal amplitude. **e**, Path of a fly in a virtual environment over 10 min. Time points were classified based on the consistency of the head direction (*ϱ*). **f**, Amplitude of PFL2 activity versus head direction for high and low *ϱ* (mean ± s.e.m. across flies, n=33 flies), with model predictions. **g**, Right-left difference in PFL3 activity for high and low *ϱ* (mean of z-scored values ± s.e.m. across flies, n=23 flies), with model predictions. **h**, Path of one fly during two 10-min epochs with high and low *ϱ*. **i**, Left: spatial profile of PFL2 activity during these same epochs. Data are divided into 4 bins based on the fly’s directional error, relative to its current goal. Right: model fits, used to infer values of *A*. **j**, Fitted goal amplitude (*A*) versus *ϱ* (mean ± s.e.m. across flies, n=33 flies). **k**, Total PFL2 activity after 180° jumps and right-left difference in PFL3 activity after 90° jumps (mean ± s.e.m. across flies, n=16 and 16 flies). Trials are classified based on whether the fly corrected its head direction after the jump. The plot on the right combines data from +90° jumps (ΣPFL3>R - ΣPFL3>L) and −90° jumps (ΣPFL3>L - ΣPFL3>R). See also Extended Data Fig. 3.

Based on this anatomy, we model the influence of the direct pathway as the right-left difference in PFL3 activity. Meanwhile, we model the influence of the indirect pathway as the total activity of PFL2, multiplied by the right-left difference in PFL3. The fly’s rotational velocity is the summed influence of both pathways. In this model, the direct pathway mainly influences steering around the goal, whereas the indirect pathway boosts steering around the anti-goal (Fig. 4b).

This model allows us to predict how head direction evolves over time. In particular, it allows us to predict how navigation should be influenced by the amplitude of the goal input signal (*A*) when we challenge the steering system with random perturbations (Fig. 4c). It shows that increasing *A* should improve the consistency of the fly’s head direction in the face of these perturbations (Fig. 4d).

To test this idea, we measured head direction consistency (*ϱ*) in a sliding 30-sec time window (Fig. 4e). We initially divided behavior into two categories, high *ϱ* (very consistent head direction) or low *ϱ* (less consistent; Fig. 4f). At each time point, we inferred the fly’s goal from its time-averaged head direction. During epochs of very consistent head direction (high *ϱ*), we found that the amplitude of PFL2 activity was strongly tuned to the fly’s directional error, with a maximal amplitude around the anti-goal (Fig. 4f). Similarly, the right-left difference in PFL3 activity was also strongly tuned to directional error (Fig. 4g). However, during epochs when the head direction was less consistent, PFL2 and PFL3 activity was much less tuned to head direction (Fig. 4f,g). These results match the predictions of our model when *A* and *ϱ* are high versus low (Fig. 4f,g). We then asked if we could infer the fly’s goal amplitude from its neural activity. Specifically, we used our network model to fit *A* at each time point in our PFL2 imaging data, and we compared these fitted values to the consistency of the fly’s head direction (*ϱ*) at each time point. We found a graded relationship between *A* and *ϱ* (Fig. 4j), as predicted (Fig. 4d). Together, these results support the idea that variations in goal amplitude (*A*) can produce variations in the organisms’ behavioral commitment to its goal (*ϱ*).

We also found evidence of behavioral variations in the responses to jumps of the virtual environment. We divided jump-trials into two categories, based on whether or not the fly made a corrective turn just after the jump. We found that, on trials when the fly turned after a 90° jump, there was a larger right-left difference in PFL3 activity. Meanwhile, on trials where the fly turned after a 180° jump, there was a larger increase in total PFL2 activity (Fig. 4k; Extended Data Fig. 3b). This result supports the idea that the brain can adjust the vigor of goal-oriented navigation behavior by varying the goal input to PFL2&3 cells.

### Navigation dynamics at cellular resolution

Thus far, we have focused testing the predictions of our model using population calcium imaging and behavioral measurements. However, we can also make predictions at the level of single-cell physiology. First, connectome data imply that most of the head direction input to PFL cells comes from inhibitory inputs (Δ7 cells) in the protocerebral bridge^5,24,30^ (Fig 5a). Whenever head direction changes, we should observe a systematic change in inhibitory synaptic input, with maximal disinhibition at the cell’s preferred head direction (Fig. 5a).

**Figure 5:**
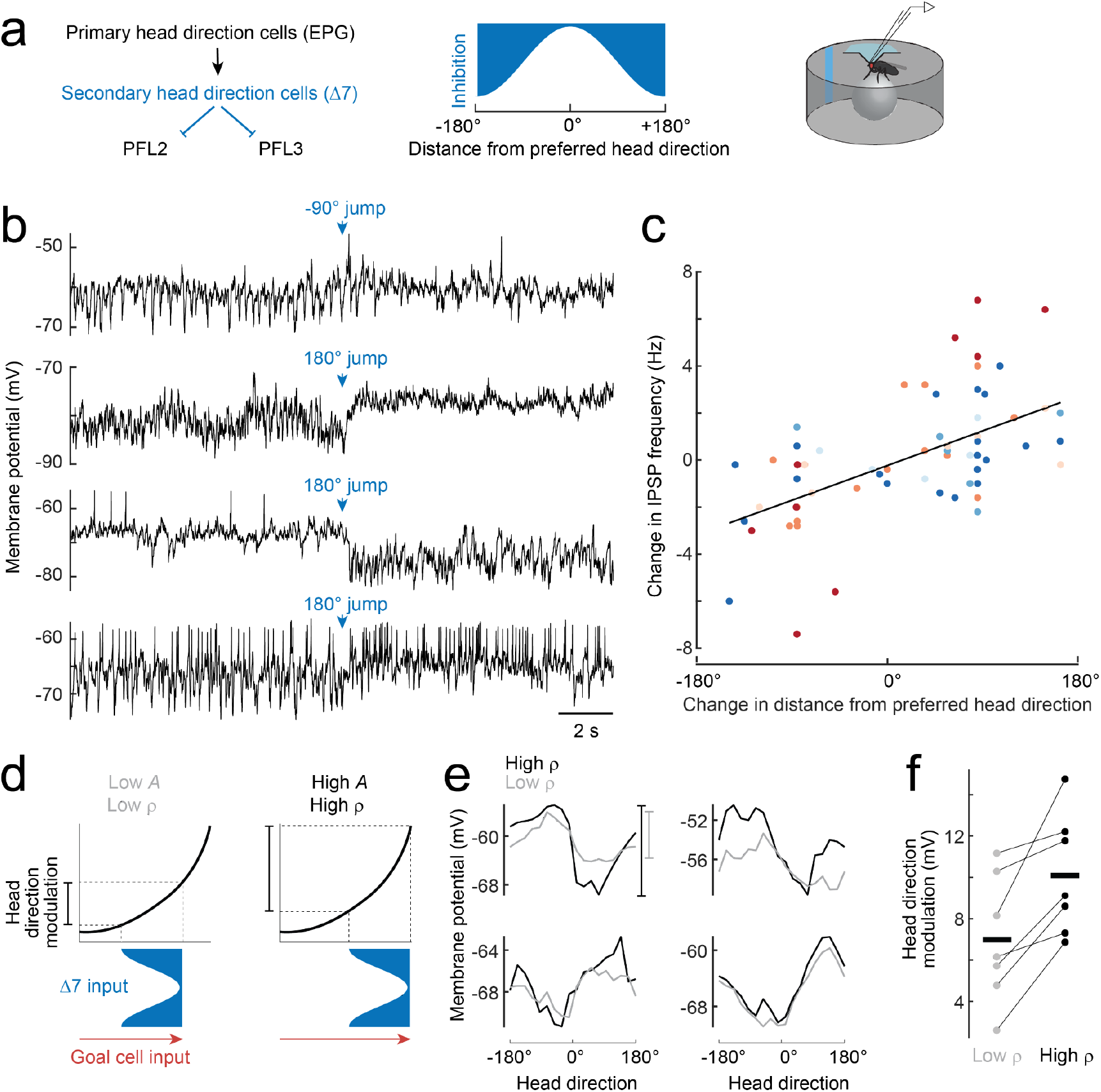
Navigation dynamics at cellular resolution. **a**, Model: Each PFL cell is predicted to receive synaptic inhibition from Δ7 cells that varies sinusoidally with head direction. We can test this prediction via whole-cell recordings from PFL cells. **b**, Whole-cell recordings from four example PFL2 and PFL3 cells show large and sustained changes in IPSP frequency when the virtual environment is rotated around the fly, emulating a change in head direction. **c**, Change in IPSP frequency versus change in head direction (relative to the cell’s preferred direction, mean ± s.e.m. across cells, n=4 PFL3 and 3 PFL2 cells in 7 flies, Pearson’s R=0.57). Each color denotes a different cell. The effect of head direction is highly significant (p=7×10^−5^, 2-way ANOVA). This analysis only uses data collected when the fly was standing still, because this makes individual IPSPs more clearly detectable (Extended Data Fig.9). **d**, Model: at the level of an individual PFL cell, goal cell input provides an excitatory boost that does not change with head direction. A larger goal amplitude *A* produces a larger boost. This boost moves the cell to a steeper part of its nonlinear input-output function. As a result, the same Δ7 input produces a larger modulation of the cell’s membrane with head direction changes. In our model, this is not due to any direct effect of *A* on Δ7 input; rather, it reflects an intrinsic nonlinearity at the level of the PFL cell. In our model, high *A* also produces high *ϱ*. **e**, Mean membrane potential versus head direction during time points when the fly was walking very straight (high *ϱ*) or less straight (low *ϱ*), for 4 example cells, all recorded in different flies. **f**, Head direction modulation (max-min) is significantly higher during epochs of goal-oriented navigation versus exploration (p =0.016, paired Wilcoxon signed rank test, n=4 PFL3 and 3 PFL2 cells in 7 flies).

Indeed, in whole-cell patch-clamp recordings, we found that this prediction was correct. Individual PFL2&3 cells exhibited approximately sinusoidal tuning to head direction, and they were continuously bombarded by large inhibitory postsynaptic potentials (IPSPs) whose frequency varied inversely with the cell’s firing rate. IPSP frequency depended strongly on the fly’s head direction, particularly when we jumped the visual environment around the fly, simulating an abrupt change in head direction (Fig. 5b, Extended Data Fig. 7). The jump often produced an abrupt and sustained increase or decrease in IPSP frequency, with a lower rate of IPSPs around the cell’s preferred head direction (Fig. 5c). This result supports the prediction that the head direction input to these cells arises mainly from inhibitory Δ7 cells.

Second, our network model predicts that increasing goal amplitude (*A*) should increase the range of PFL2&3 activity fluctuations. This is because increasing goal amplitude (*A*) produces more excitation onto PFL2&3 cells; this shifts the net input to every PFL2&3 cell to a more positive value, placing it on a steeper part of its nonlinear input-output function (Fig. 5d). As a result, the cell should be more sensitive to changes in synaptic input from the head direction system. If this prediction is correct, it would provide direct evidence for the nonlinear step in our model.

To test this prediction, we examined the fly’s behavior during our whole-cell recordings, to infer the fly’s goal amplitude. As before, we classified each time point as goal-oriented navigation or exploration, based on the consistency of the fly’s head direction around that time (*ϱ*). During epochs of goal-oriented navigation, when the goal amplitude (*A*) is likely to be high, we found that PFL2&3 membrane potential was a relatively steep function of head direction. Conversely, during epochs of exploration, when *A* is likely to be low, PFL2&3 membrane potential was only shallowly dependent on head direction (Fig. 5e,f). Together, these results corroborate the idea that head direction inputs and goal inputs are combined nonlinearly in PFL2&3 cells. Moreover, these results argue that the nonlinearity operates at the level of voltage-gated conductances that control the sensitivity of the membrane potential to synaptic inputs. The goal amplitude *A* influences the steepness of this function, likely by simply shifting each PFL2 or 3 cell to a more depolarized voltage.

## Discussion

The brain’s maps of space are allocentric, meaning that they encode the organism’s location relative to objects in the world. Conversely, motor commands are intrinsically egocentric. Thus, during navigation, there must be a transfer of information from allocentric to egocentric coordinates. Moreover, the mapping from spatial coordinates to cells is flexible: for example, a given head direction may be represented by specific cells at one time, but by other cells at a later time^31^. This phenomenon of representational drift^32–34^ is a natural consequence of the fact that the brain’s spatial maps are plastic and continuously changing^35–38^. Finally, spatial goals are often contingent on circumstances. For example, an organism may need to head away from its nest to find food, but then reverse its goal to head back to its nest once food has been found^39^. In short, the brain must store information about where it should go, referencing this information to a constantly evolving map of space, and then retrieve that information for subsequent action control.

Here we describe a network architecture that allows head direction cells to control steering, based on an internal goal. Our results show how the allocentric head direction map can be transformed into egocentric steering commands. This network creates three shifted copies of the head direction representation, with each copy instantiated in a different population of cells; these three copies tile the space of possible compass directions. All three populations also share an allocentric goal input. Each population detects the overlap between head direction input and goal input, and together, their output produces an appropriate egocentric steering drive.

It seems likely that there are many sets of “goal cells” in the brain^9–11^. There are dozens of cell types in the fan-shaped body that provide strong direct input to PFL2 and PFL3 cells, and many of these cell types have the appropriate geometrical organization to represent a goal as a spatial sinusoid^5^. In principle, goals could be stored as spatial patterns of persistent activity or synaptic weights. The signal to initialize these patterns may be gated by input from the mushroom body to the fan-shaped body^5,11,29,40–43^ based on associations formed between head direction and reward or punishment^10,13,44^. Goal signals may also be integrated over long time periods; indeed, the representation of the path integral can be treated as a kind of a goal^9,45^, and it could be stored in a similar format, i.e. a spatial sinusoid.

The network architecture we have described also suggests a solution to the challenge posed by representational drift. Specifically, as the phase of the head direction representation drifts over time during spatial learning^35–37^, the goal representation simply needs to shift in concert. The same learning rule that initialized the goal representation could update that representation every time a reward or punishment is repeated, keeping the goal aligned with the current coordinate frame of the head direction system. As a result, motor commands would remain aligned to the allocentric goal; this might explain why representational drift is less obvious in cells more strongly correlated with motor performance^46^.

Importantly, our results also have implications for the logic of motor control. If a motor system does not update rapidly, its performance will lag; conversely, if it updates quickly with a gain that is too high, its performance will overshoot. Thus, motor feedback loops need to be adjusted to match the current needs of the system, a concept known as adaptive control^47^. We propose that PFL2 cells essentially provide adaptive control of steering, by increasing the gain of the directional signal, from PFL3 cells, when error is large; conversely, this allows gain to be low when error is small, thereby avoiding overshooting. In our model, the adaptive control exerted by PFL2 cells occurs only in the “indirect” pathway to the motor system. This indirect pathway is augmented by a direct pathway from PFL3 cells to descending neurons; the direct pathway ensures that steering gain never drops too low. In the future, it will be interesting to investigate the predictions of this model from a mechanistic perspective, focusing on the intermediate cells in this pathway (DNa03). We note that a recent study^10^ proposed a conceptually similar role for PFL2 cells, but framed in terms of statistical drift during reinforcement learning.

In summary, our results reveal how the sense of direction can be used to generate adaptive locomotor commands. Our conclusions generate testable new predictions for how multiple goals could be stored in memory, adjusted for salience, and retrieved on demand. Because the basic problems of navigation are fundamental problems of geometry and information retrieval, the solutions we describe here may have general relevance for other systems.

## Author contributions

E.A.W. performed all the experiments and analyzed the data. E.K., P.M.D., and J.L. contributed to the identification and characterization of split-Gal4 lines. L.H., B.M., S.D., E.A.W., and R.I.W. contributed to conceptualization and modeling.

## Acknowledgements

We thank Nils Eckstein, Alexander S. Bates, Andrew Champion, Gregory S.X.E. Jefferis, and Jan Funke for early access to neurotransmitter prediction results for the hemibrain connectome. Michael Dickinson and the Research Instrumentation Core at Harvard Medical School provided hardware/software assistance. We are grateful to Michael Dickinson and Aleksandr Rayshubskiy for helpful conversations, and to Stephen Holtz, Matt Collie, and Noah Pettit for comments on the manuscript. This work was supported by NIH grant U19NS104655 (to R.I.W. and S.D.). R.I.W. is an HHMI Investigator.

---

**Extended Data Fig. 1:**
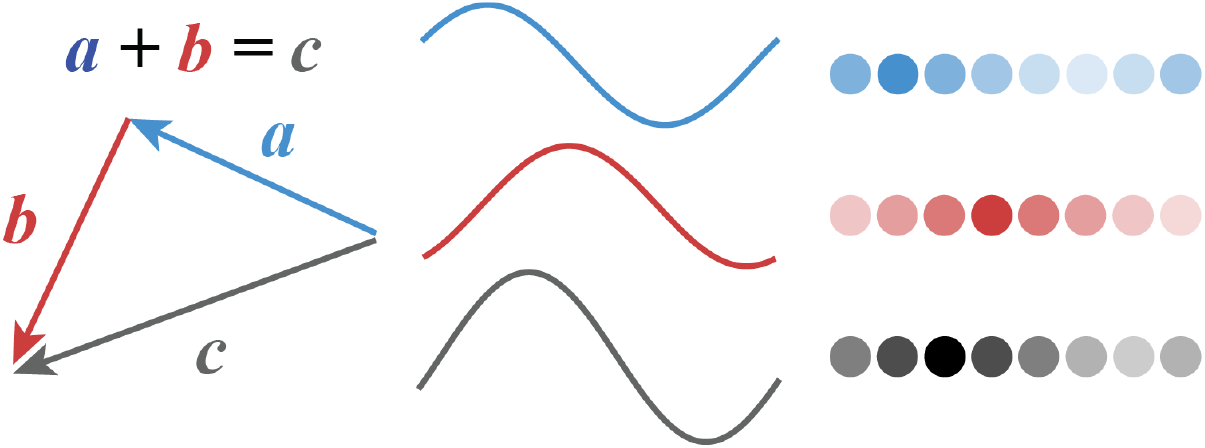
Representing vectors as sinusoids. Any vector can be represented as a sinusoidal function whose amplitude represents the magnitude of the vector, and whose phase represents the angle of the vector. Although it is convenient to represent these sinusoids as continuous functions, they can also be discretized into spatial activity patterns over populations of discrete neurons^15,16^. Adding these sinusoids pointwise is equivalent to performing vector addition.

**Extended Data Fig. 2:**
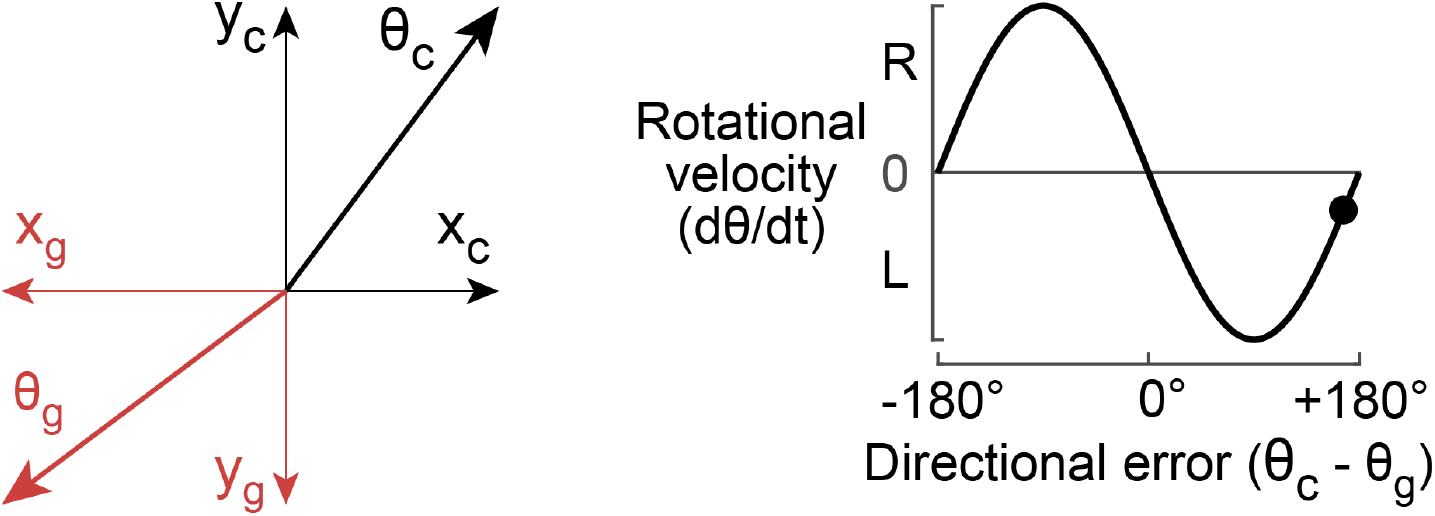
Resolver servomechanisms. A resolver measures the current angle of some object (θ_c_, e.g., the angular position of a shaft) and resolves that angle into its Cartesian components (x_c_ and y_c_). The goal angle (θ_g_) is similarly resolved into its Cartesian components (x_g_ and y_g_). These components are then cross-multiplied, and their difference is taken as a rotational velocity command:

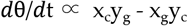 The schematic above treats positive values of *ϱ*θ/*ϱ*t as clockwise rotations, and negative values of of *ϱ*θ/*ϱ*t as counter-clockwise rotations. In this example, the current angle is rotated clockwise relative to the goal, meaning a positive directional error. This drives a counterclockwise rotation. But because the directional error at this point (•) is almost 180°, the rotational velocity command is relatively small. When θ_c_ = θ_g_ + 180°, the rotational velocity command is 0; this requires additional measures to prevent “false nulling” of the servomechanism. Mittelstaedt suggested that a similar process might be implemented in the brain’s navigation centers to control an organism’s heading, and thus its path through the environment; this is known as “Mittelstaedt ‘s bicomponent model” of steering control^7^. We show that PFL3 neurons are conceptually somewhat similar to a resolver servomechanism: they produce right-left shifted copies of the original angle representation, and they compare these copies to a goal angle. These quantities are then subtracted to produce a signal that varies sinusoidally with the system’s directional error.

**Extended Data Fig. 3:**
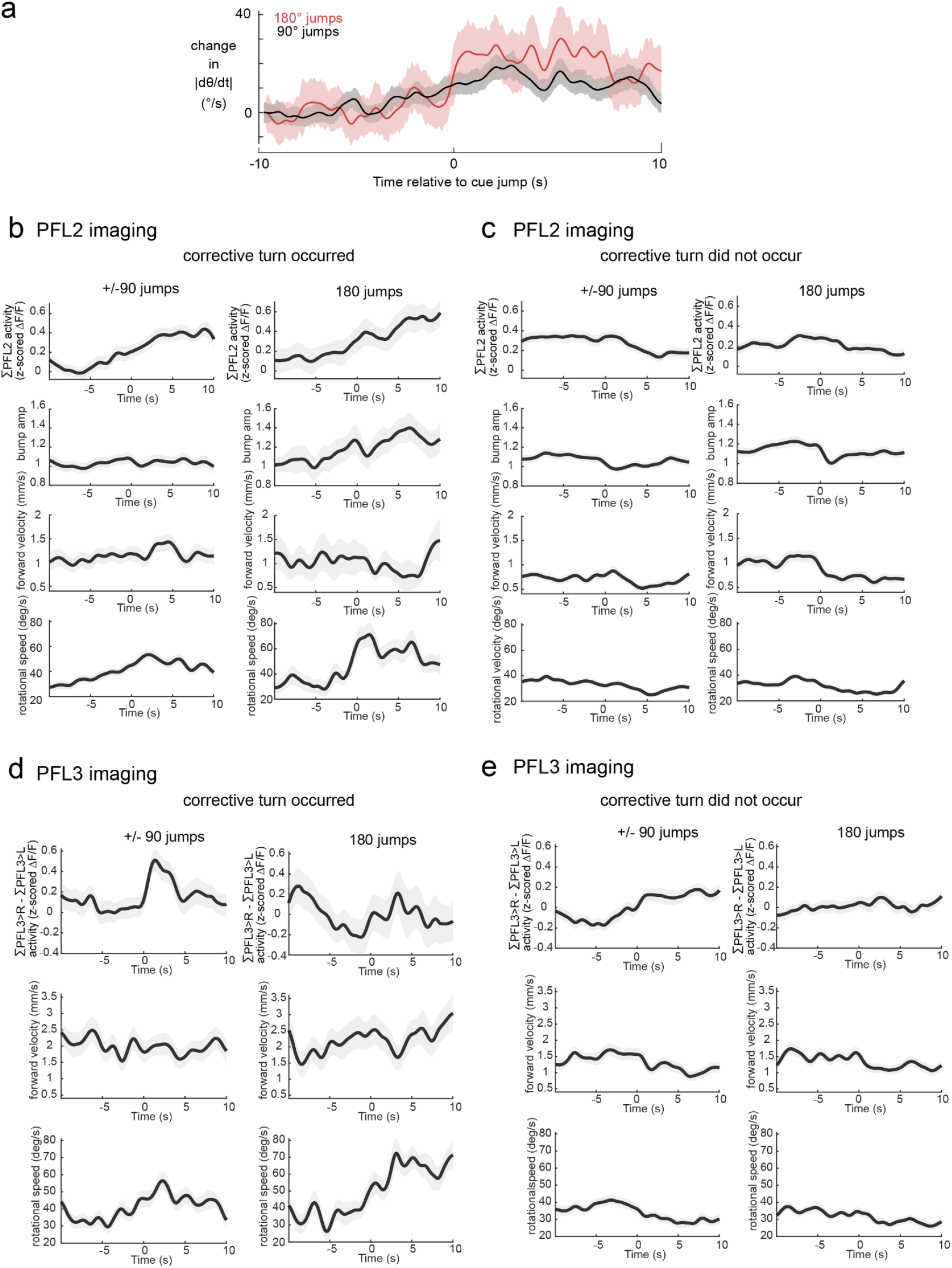
Changes in neural activity and behavior for 90°, − 90°, and 180° cue jumps. **a**, Jumping the environment by 90° and 180° produces a similar change in rotational speed. Bands are s.e.m. across flies (n=46 flies). **b**. The leftmost column of panels show changes in PFL2 activity and behavior following +/- 90° cue jumps in which the jump was categorized as having been corrected, such that the fly had been walking relatively straight prior to the jump and within the following 10 seconds had returned it to a position within +/- 30° of its original position (mean ± s.e.m, n=31 flies). The right column shows the same but for 180° jumps (mean ± s.e.m, n=16 flies). **c**. The same as (a) except for jumps that were categorized as uncorrected (mean ± s.e.m, n=33 flies). The right column shows the same but for 180° jumps (mean ± s.e.m, n=29 flies). **c**. the same as (a) except for PFL3 activity (mean ± s.e.m, n=16 flies). The right column shows the same but for 180° jumps (mean ± s.e.m, n=11 flies). **d**. The same as (b) except for PFL3 activity (mean ± s.e.m, n=23 flies). The right column shows the same but for 180° jumps (mean ± s.e.m, n=23 flies).

**Extended Data Fig. 4:**
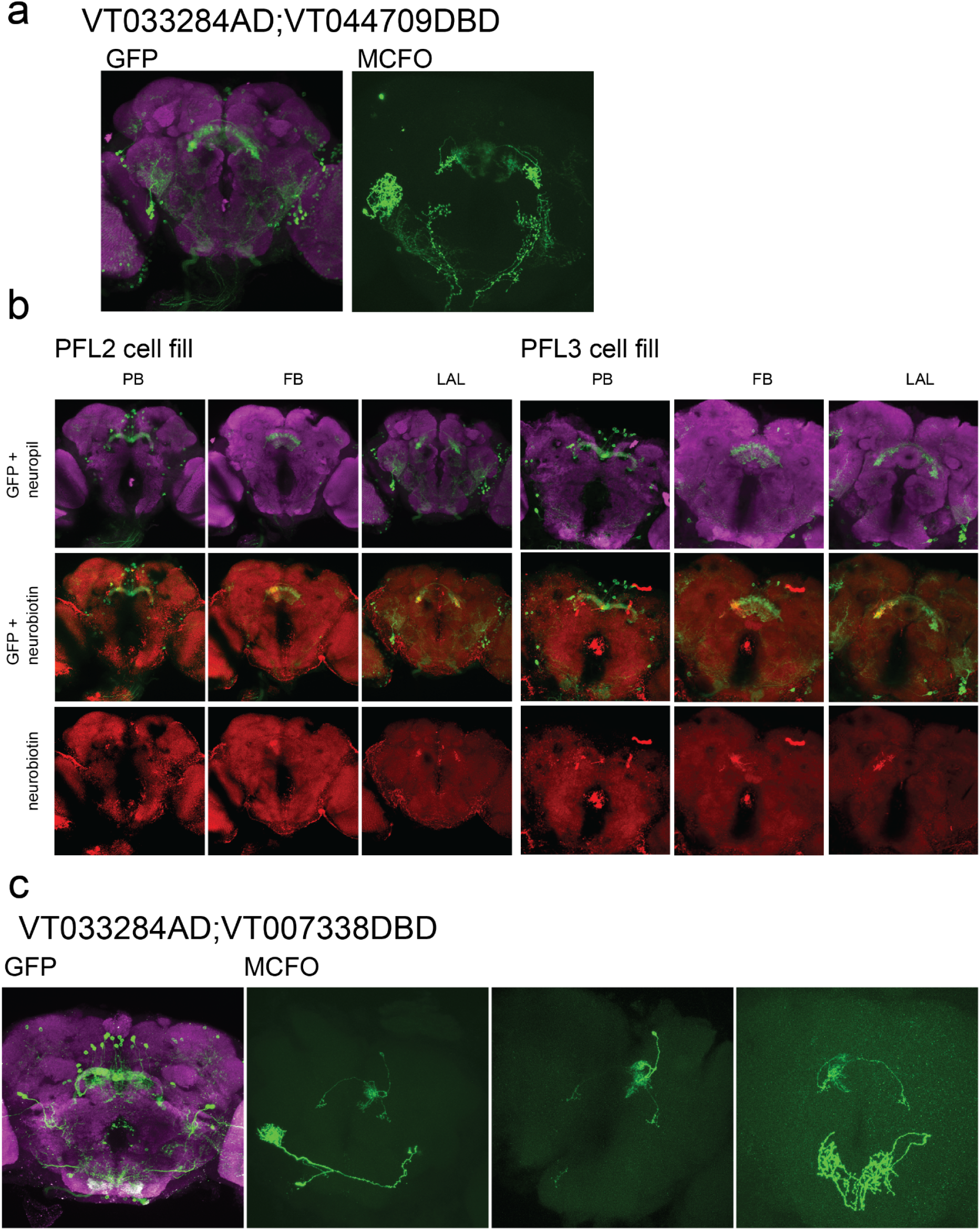
Example images for mixed and PFL2 split-Gal4 lines. Left panel shows a max z-projection of the mixed PFL2/3 split-Gal4 line expressing GFP. Right panel shows a max z-projection from an MCFO clone with 3 PFL3 neurons (PFL3 identity confirmed by counting axons) along with unidentified neurons outside of the central complex. Across 7 clones we counted 7 PFL3 neurons and 4 PFL2 neurons, no expression from other cell types was observed in the region of the lateral accessory lobes where PFL2 and PFL3 axon terminals are found. **b**. Example cell fills obtained from electrophysiology experiments. The left 3 columns show an example PFL2 cell fill while the right 3 columns show an example PFL3 cell fill. It was common for the axonal arbors to exhibit greater fluorescence compared to the dendritic arbors in both PFL3 and PFL2 fills. **c**. Left panel shows a max z-projection of the PFL2 split-Gal4 line expressing GFP. The right three panels each show a max z-projection of a different MCFO clone each with one PFL2 neuron. The left and rightmost of these panels also show an unidentified neuron ventral to the central complex. Across 13 clones we counted 15 PFL2 neurons, and 0 other cell types in the central complex.

**Extended Data Fig. 5:**
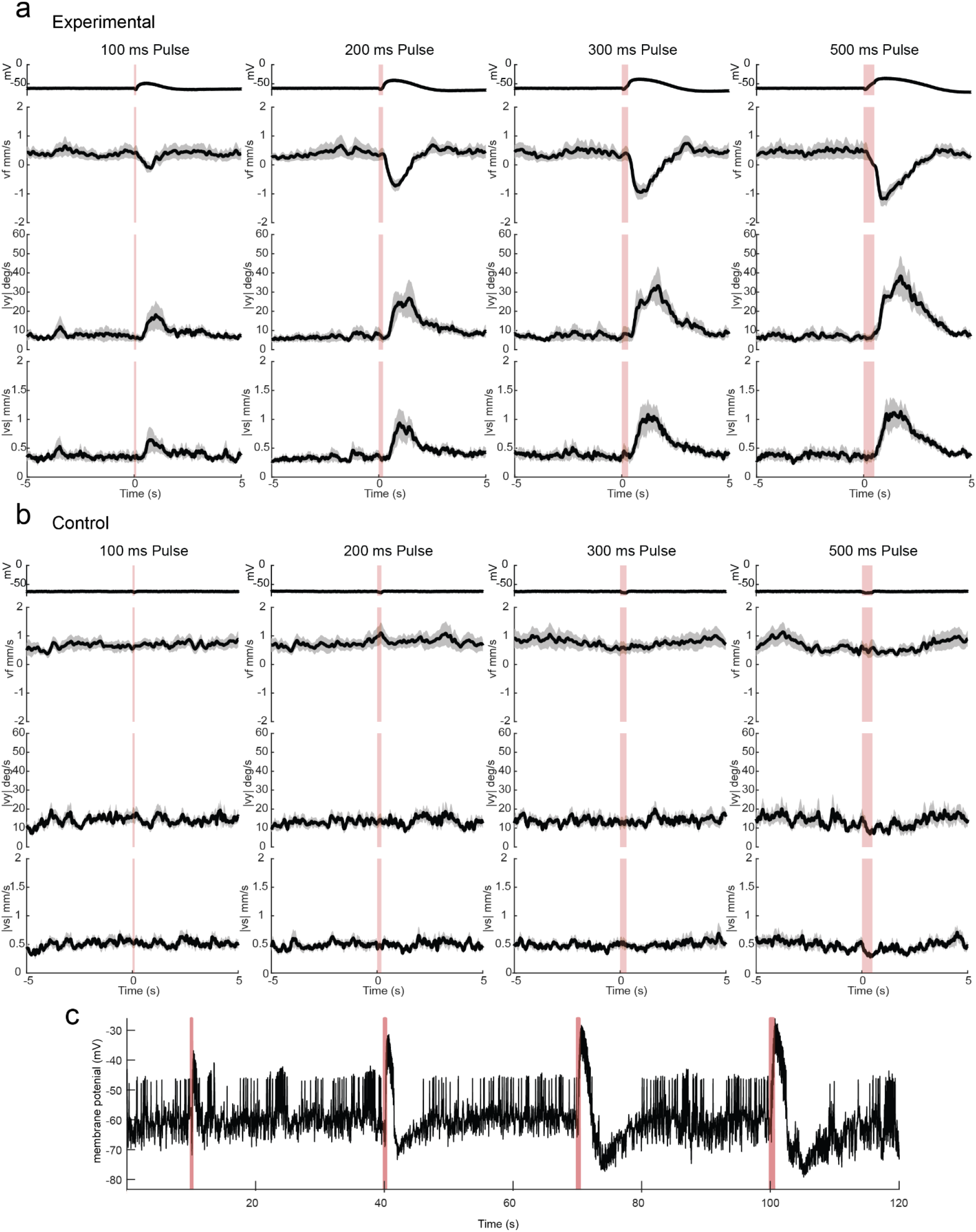
Expanded Iontophoresis data. **a**. Expanded summary data for flies where PFL2 cells expressed P2×2 showing 100ms, 200ms, 300ms, and 500ms pulses and the subsequent response in forward velocity (vf), rotational speed (|vy|), and sideways speed (|vs|) **b**. Expanded summary data for genetic controls showing 100ms, 200ms, 300ms, and 500ms pulses and the subsequent response in forward velocity (vf), rotational speed (|vy|), and sideways speed (|vs|) **c**. Example raw membrane potential trace showing the change in membrane potential following a 100ms, 200ms, 300ms, and 500ms pulse over a 120 second period.

**Extended Data Fig. 6:**
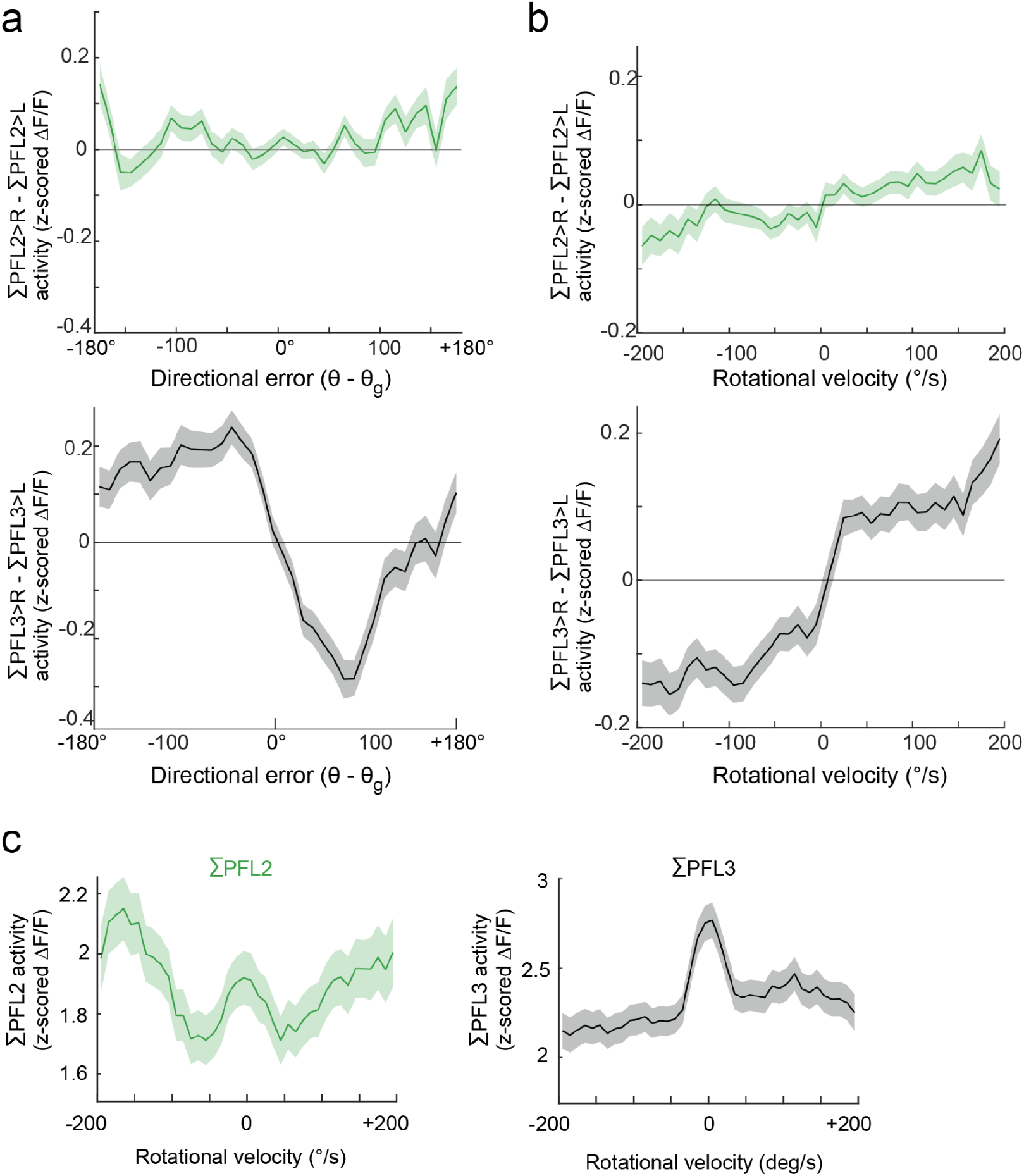
Comparison of patterns of LAL activity between the PFL2 and mixed line. **a**. Top panel: Right minus left PFL2 activity in the LAL versus directional error (mean ± s.e.m. n=33 flies). Bottom panel: the same but for PFL2+3 neurons. (mean ± s.e.m. n=23 flies) **b**. Top panel: Right minus left PFL2 activity in the LAL versus rotational velocity(mean ± s.e.m. n=33 flies). Bottom panel: the same but for PFL2+3 neurons. (mean ± s.e.m. n=23 flies) **c**. Left panel: summed PFL2 activity in the LAL, versus rotational velocity (mean ± s.e.m. n=33 flies). Right panel: the same but for PFL2+3 neurons. (mean ± s.e.m. n=23 flies).

**Extended Data Fig. 7:**
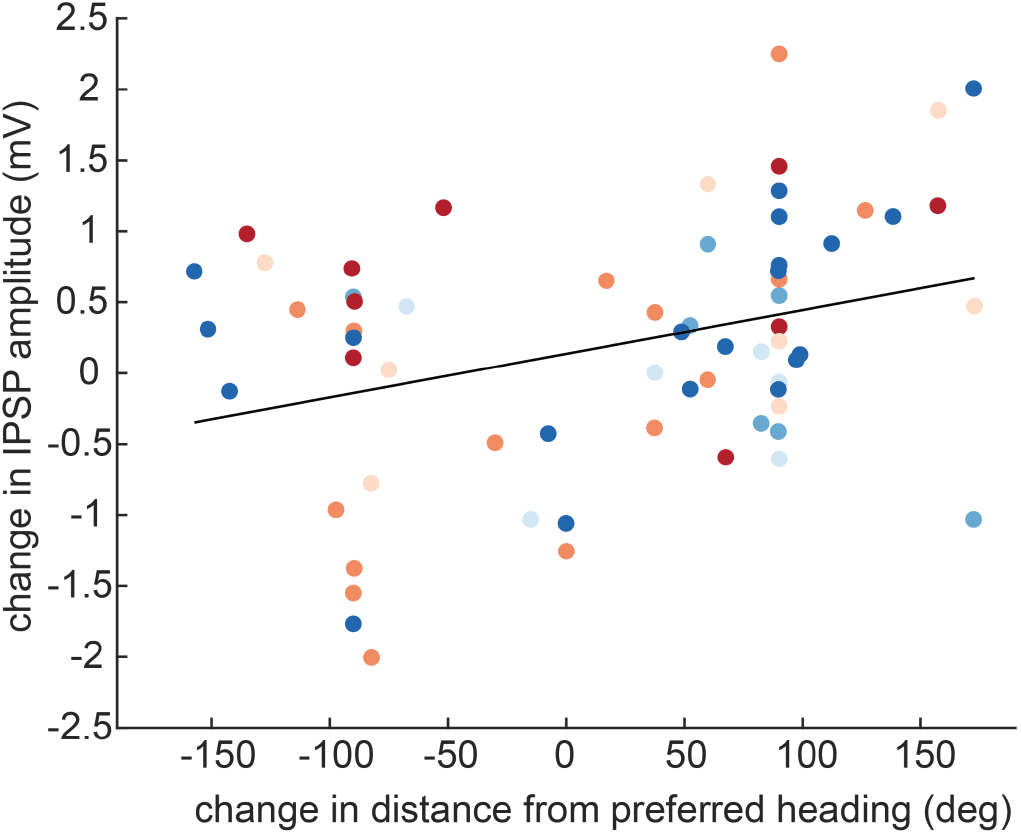
Changes in IPSP amplitude relative to distance from preferred heading. Similar to Fig 5c but now showing the change in IPSP amplitude versus change in head direction (relative to the cell’s preferred direction, mean ± s.e.m. across cells, n=4 PFL3 and 3 PFL2 cells in 7 flies, Pearson’s R=0.33). Each color denotes a different cell. The effect of head direction is not significant (p= 0.22, 2-way ANOVA). This analysis only uses data collected when the fly was standing still, because this makes individual IPSPs more clearly detectable.

**Extended Data Fig. 8:**
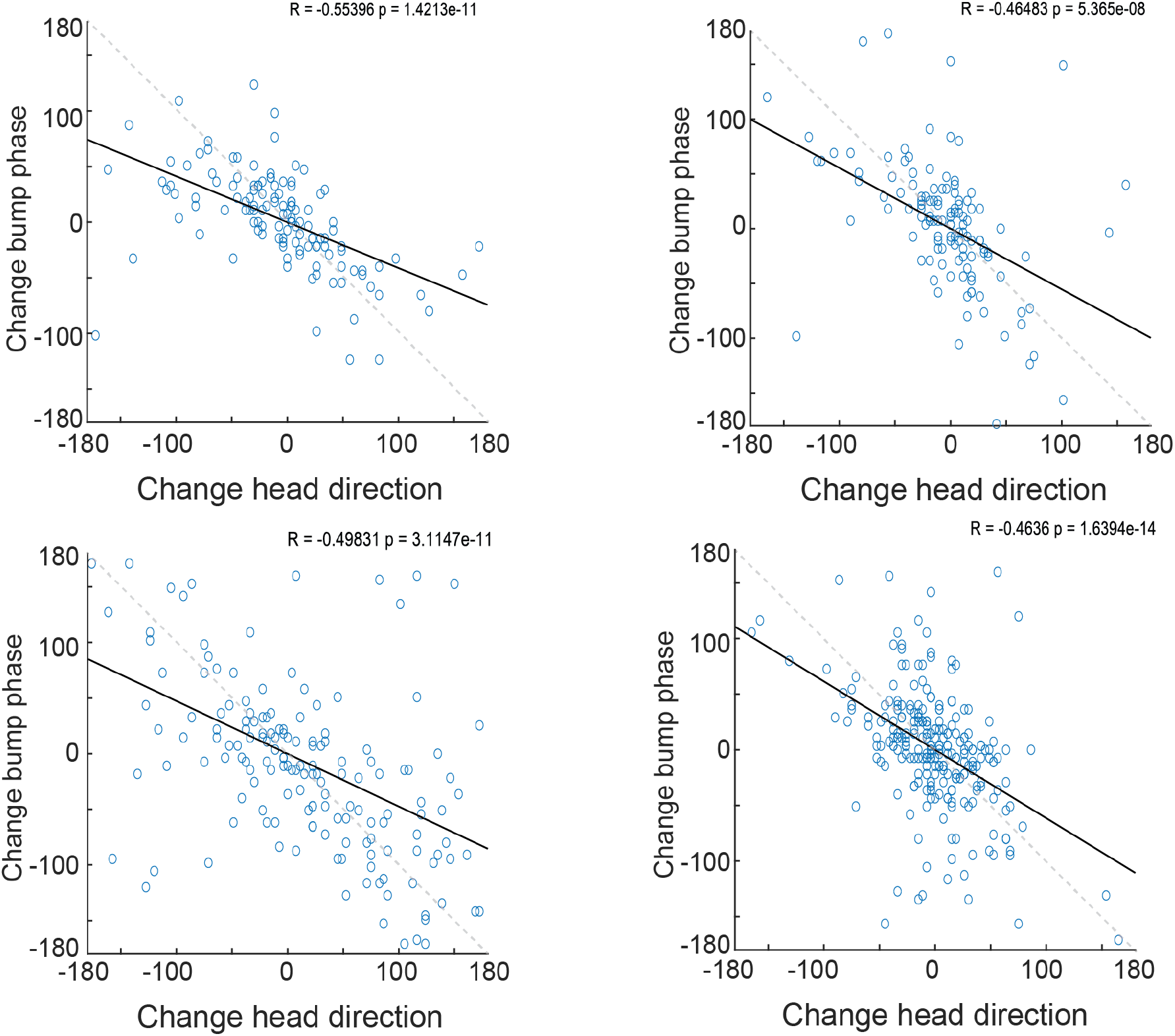
Examples of the relationship of changes in PFL2 bump phase versus changes in head direction. Change in PFL2 phase versus change in head direction for 4 more example flies, the same as shown in Fig 2d, Pearson’s R and p value calculated for the linear fit displayed on each plot. In each example as the fly’s head direction rotates clockwise the PFL2 phase rotates counterclockwise.

**Extended Data Fig. 9:**
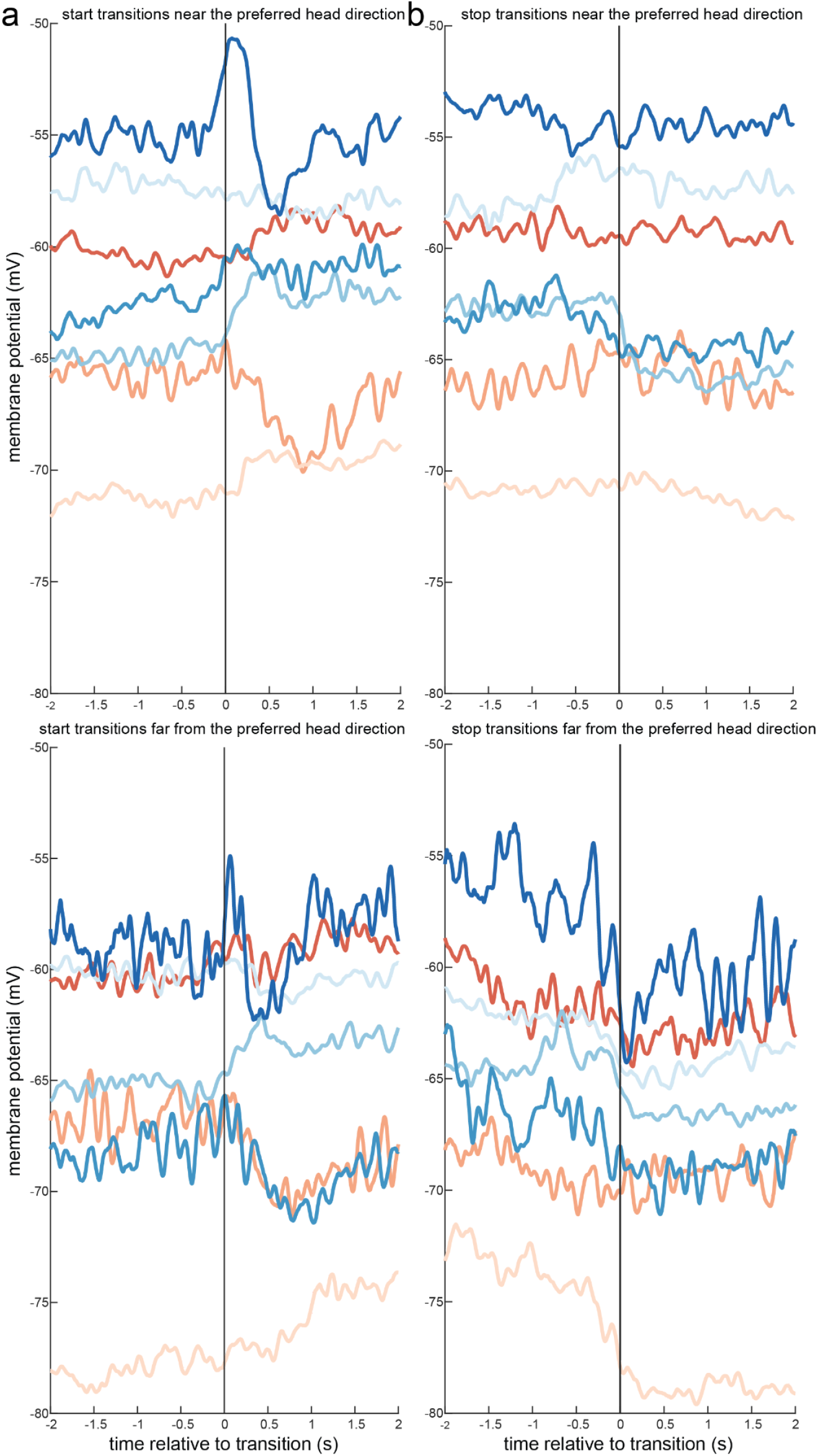
Average changes in membrane potential across 7 cells around start & stop transitions. **a**. The top panel shows the membrane potential for each of the 7 cells against time relative to a transition from no movement to movement when the fly had been facing within +/- 45° of each cell’s preferred head direction. The bottom panel shows the same except that transition occurred when the fly was facing further than +/- 135° of the cell’s preferred head direction. **b**. The same as a but now for stop transitions where the fly changed from a moving to a not moving state.

## Methods

### Flies

Unless otherwise specified, flies were raised on cornmeal-molasses food (Archon Scientific) in an incubator on a 12 h:12 h light:dark cycle at 25 °C at 50–70% relative humidity. Experimenters were not blinded to fly genotype. For iontophoresis stimulus experiments (Fig. 2a,b) flies were grouped for analysis based on genotype. Sample sizes were chosen based on conventions in our field for standard sample sizes; these sample sizes are conventionally determined on the basis of the expected magnitude of animal-to-animal variability, given published results and pilot data. All experiments used flies with at least one wild-type copy of the white (*w*) gene. Genotypes of fly stocks used in each figure are as follows.

Fig. 1:

PFL2/3 calcium imaging, w/+; P{VT033284-p65AD}attP40/20XUAS-IVS-cyRFP{VK00037}; P{y[+t7.7];w[+mC]=VT044709-GAL4.DBD}attP2/PBac{y[+t7.7] w[+mC]=20XUAS-IVS-jGCaMP7b}VK00005

PFL2 calcium imaging, w/+; P{VT033284-p65AD}attP40/20XUAS-IVS-cyRFP{VK00037}; P{y[+t7.7];P{VT007338-Gal4DBD}attP2/PBac{y[+t7.7] w[+mC]=20XUAS-IVS-jGCaMP7b}VK00005

Fig. 2:

PFL2 line iontophoresis,

w/+;P{VT033284-p65AD}attP40/P{w[+mC]=UAS-Rnor\P2rx2.L}4/;P{VT007338-Gal4DBD}attP2/20XUAS-mCD8:

:GFP {attP2}

Empty split control iontophoresis, w/+;P{y[+t7.7] w[+mC]=p65.AD.Uw}attP40/P{w[+mC]=UAS-Rnor\P2rx2.L}4;P{y[+t7.7] w[+mC]=GAL4.DBD.Uw}attP2/20XUAS-mCD8::GFP {attP2}

PFL2 calcium imaging, w/+; P{VT033284-p65AD}attP40/20XUAS-IVS-cyRFP{VK00037}; P{y[+t7.7];P{VT007338-Gal4DBD}attP2/PBac{y[+t7.7] w[+mC]=20XUAS-IVS-jGCaMP7b}VK00005

Fig. 3:

PFL2 calcium imaging, w/+; P{VT033284-p65AD}attP40/20XUAS-IVS-cyRFP{VK00037}; P{y[+t7.7];P{VT007338-Gal4DBD}attP2/PBac{y[+t7.7] w[+mC]=20XUAS-IVS-jGCaMP7b}VK00005

PFL2/3 calcium imaging, w/+; P{VT033284-p65AD}attP40/20XUAS-IVS-cyRFP{VK00037}; P{y[+t7.7];w[+mC]=VT044709-GAL4.DBD}attP2/PBac{y[+t7.7] w[+mC]=20XUAS-IVS-jGCaMP7b}VK00005

Fig. 4:

PFL2 calcium imaging, w/+; P{VT033284-p65AD}attP40/20XUAS-IVS-cyRFP{VK00037}; P{y[+t7.7];P{VT007338-Gal4DBD}attP2/PBac{y[+t7.7] w[+mC]=20XUAS-IVS-jGCaMP7b}VK00005

PFL2/3 calcium imaging, w/+; P{VT033284-p65AD}attP40/20XUAS-IVS-cyRFP{VK00037}; P{y[+t7.7];w[+mC]=VT044709-GAL4.DBD}attP2/PBac{y[+t7.7] w[+mC]=20XUAS-IVS-jGCaMP7b}VK00005;

Fig. 5:

w/+; P{VT033284-p65AD}attP40/P{20XUAS-IVS-mCD8::GFP}attP40;P{y[+t7.7] w[+mC]=VT044709-GAL4.DBD}attP2/+

Extended data figure 3:

PFL2 calcium imaging, w/+; P{VT033284-p65AD}attP40/20XUAS-IVS-cyRFP{VK00037}; P{y[+t7.7];P{VT007338-Gal4DBD}attP2/PBac{y[+t7.7] w[+mC]=20XUAS-IVS-jGCaMP7b}VK00005

PFL2/3 calcium imaging, w/+; P{VT033284-p65AD}attP40/20XUAS-IVS-cyRFP{VK00037}; P{y[+t7.7];w[+mC]=VT044709-GAL4.DBD}attP2/PBac{y[+t7.7] w[+mC]=20XUAS-IVS-jGCaMP7b}VK00005

Extended data figure 4:

PFL2/3 line: w/+; P{VT033284-p65AD}attP40/P{20XUAS-IVS-mCD8::GFP}attP40;P{y[+t7.7] w[+mC]=VT044709-GAL4.DBD}attP2/+

PFL2 line: w/+; P{VT033284-p65AD}attP40/P{20XUAS-IVS-mCD8::GFP}attP40; P{y[+t7.7];P{VT007338-Gal4DBD}attP2/+

Extended data figure 5:

PFL2 line iontophoresis,

w/+;P{VT033284-p65AD}attP40/P{w[+mC]=UAS-Rnor\P2rx2.L}4/;P{VT007338-Gal4DBD}attP2/20XUAS-mCD8:

:GFP {attP2}

Empty split control iontophoresis, w/+;P{y[+t7.7] w[+mC]=p65.AD.Uw}attP40/P{w[+mC]=UAS-Rnor\P2rx2.L}4;P{y[+t7.7] w[+mC]=GAL4.DBD.Uw}attP2/20XUAS-mCD8::GFP {attP2}

Extended data figure 6:

PFL2 calcium imaging, w/+; P{VT033284-p65AD}attP40/20XUAS-IVS-cyRFP{VK00037}; P{y[+t7.7];P{VT007338-Gal4DBD}attP2/PBac{y[+t7.7] w[+mC]=20XUAS-IVS-jGCaMP7b}VK00005

PFL2/3 calcium imaging, w/+; P{VT033284-p65AD}attP40/20XUAS-IVS-cyRFP{VK00037}; P{y[+t7.7];w[+mC]=VT044709-GAL4.DBD}attP2/PBac{y[+t7.7] w[+mC]=20XUAS-IVS-jGCaMP7b}VK00005

Extended data figure 7:

w/+; P{VT033284-p65AD}attP40/P{20XUAS-IVS-mCD8::GFP}attP40;P{y[+t7.7] w[+mC]=VT044709-GAL4.DBD}attP2/+

Extended data figure 8:

w/+; P{VT033284-p65AD}attP40/20XUAS-IVS-cyRFP{VK00037};

P{y[+t7.7];P{VT007338-Gal4DBD}attP2/PBac{y[+t7.7] w[+mC]=20XUAS-IVS-jGCaMP7b}VK00005

Extended data figure 9:

w/+; P{VT033284-p65AD}attP40/P{20XUAS-IVS-mCD8::GFP}attP40;P{y[+t7.7] w[+mC]=VT044709-GAL4.DBD}attP2/+

### Origins of transgenic stocks

The following stocks were obtained from the Bloomington Drosophila Stock Center (BDSC) and published as follows: P{y[+t7.7]w[+mC]=VT044709-GAL4.DBD}attP2 (BDSC_75555)^59^, P{y[+t7.7] w[+mC]=p65.AD.Uw}attP40; P{y[+t7.7] w[+mC]=GAL4.DBD.Uw}attP2 (BDSC_79603), P{w[+mC]=UAS-Rnor\P2rx2.L}4/CyO (BDSC_91223)^60^

The following stocks were obtained from WellGenetics: w[1118];P{VT007338-p65ADZp}attP40/CyO;+ (SWG9178/A), w[1118];P{VT033284-p65AD}attP40/CyO;+ (A/SWG8077). Using these lines we constructed a split-Gal4 line that targets PFL2 & 3 neurons in the lateral accessory lobes (LAL), +;P{VT033284-p65AD}attP40;P{y[+t7.7] w[+mC]=VT044709-GAL4.DBD}attP2. We validated the expression of this line using immunohistochemical anti-GFP staining, and also using Multi-Color-Flip-Out (MCFO) to visualize single-cell morphologies. This line has significant non-specific expression throughout the brain but has been determined to be specific for PFL2/3 in the lateral accessory lobes. We also constructed a split-Gal4 line to target PFL2 neurons, +; P{VT033284-p65AD}attP40; P{y[+t7.7];P{VT007338-Gal4DBD}attP2. We validated the expression of this line using immunohistochemical anti-GFP staining, and also using Multi-Color-Flip-Out (MCFO) to visualize single-cell morphologies. This line exhibits expression in various peripheral neurons but is clean for PFL2 neurons within the central complex. Specifically the protocerebral bridge, fan-shaped body, and lateral accessory lobes.

### Fly preparation and dissection

Flies used for all experiments were isolated the day prior to the experiment by single-housing on molasses food. For calcium imaging experiments we used female flies 20-72 hours post-eclosion. For electrophysiology experiments, including the iontophoresis experiments, we used female flies 16-30 hours post-eclosion. No circadian restriction was imposed for the time of experiments.

Manual dissections in preparation for experiments were as follows. Flies were briefly cold-anesthetized and inserted using fine forceps (Fine Science Tools) into a custom platform machined from black Delrin (Autotiv or Protolabs). The platform was shaped like an inverted pyramid to minimize optical occlusion. The head was pitched slightly forward so that the posterior surface was more accessible to the microscope objective. The wings were removed, then the fly head and thorax were secured to the holder using UV-curable glue (Loctite AA 3972) with a brief pulse of ultraviolet light (LED-200, Electro-Lite Co). To remove large brain movements, the proboscis was glued in place using a small amount of the same UV-curable glue. Using fine forceps in extracellular *Drosophila* saline, a window was opened in the head cuticle, and tracheoles and fat were removed to expose the brain. To further reduce brain movement, muscle 16 was stretched by gently tugging the esophagus, or else it was removed by clipping the muscle anteriorly. For electrophysiology and iontophoresis experiments only, the perineural sheath was minimally removed with fine forceps over the brain region of interest. For all experiments, saline was continuously superfused over the brain. *Drosophila* extracellular saline composition was: 103 mM NaCl, 3 mM KCl, 5 mM TES, 8 mM trehalose, 10 mM glucose, 26 mM NaHCO3, 1 mM NaH2PO4, 1.5 mM CaCl2, and 4 mM MgCl2 (osmolarity 270-275 mOsm). Saline was oxygenated by bubbling with carbogen (95% O2 / 5% CO2), and reached a final pH of ∼7.3.

### Two-photon calcium imaging

We used a two-photon microscope equipped with a galvo-galvo-resonant scanhead (Thorlabs Bergamo II GGR) and 25× 1.10 N.A. objective (Nikon CFI APO LWD; Thorlabs, WDN25X-APO-MP). For volumetric imaging we used a fast piezoelectric objective scanner (Thorlabs PFM450E). To excite GCaMP we used a wavelength-tunable femtosecond laser with dispersion compensation (Mai Tai DeepSee, Spectra Physics) set to 920 nm. GCaMP fluorescence signals were collected using GaAsP PMTs (PMT2100, Thorlabs) through a 405/488 nm bandpass filter (Thorlabs). All image acquisition and microscope control was conducted in Matlab 2021a (MathWorks Inc) using ScanImage 2021 Premium with vDAQ hardware (Vidrio Technologies LLC, Ashburn, VA), and custom Matlab scripts for further experimental control. The imaging region for fan-shaped body (FB) and protocerebral bridge (PB) experiments was 150×250 pixels, for lateral accessory lobe (LAL) experiments it was 150×400 pixels, with 10-12 slices in the z-axis for each volume (4 μm per slice), resulting in 6-8 Hz volumetric scanning rate. For experiments using the selective PFL2 split-Gal4 line, we imaged in the PB, FB, or LAL for different trials. For experiments imaging the mixed PFL3+PFL2 split-Gal4 line, we only imaged in the LAL.

### Patch-clamp recordings

Patch pipettes were pulled from filamented borosilicate capillary glass (OD 1.5 mm, ID 0.86 mm; BF150-86-7.5HP, Sutter Instrument Company), using a horizontal pipette puller (P-97, Sutter Instrument Company) to a resistance range of 9–13 MΩ. Pipettes were filled with an internal solution^48^ consisting of 140 mM KOH, 140 mM aspartic acid, 1 mM KCl, 10 mM HEPES, 1 mM EGTA, 4 mM MgATP, 0.5 mM Na_3_GTP, and 15 mM neurobiotin citrate, filtered twice through a 0.22-μm PVDF filter (Millipore).

All electrophysiology experiments used a semi-custom upright microscope consisting of a motorized base (Thorlabs Cerna), with conventional collection and epifluorescence attachment (Olympus BX51), but no substage optics in order to better fit the virtual reality system. The microscope was equipped with a 40X water immersion objective (LUMPlanFLN 40XW, Olympus) and CCD Monochrome Camera (Retiga ELECTRO; 01-ELECTRO-M-14-C Teledyne) For GFP excitation and detection, we used a 100 W Hg arc lamp (Olympus U-LH100HG) and an eGFP long-pass filter cube (Olympus F-EGFP LP). The fly was illuminated from below using a fiber optic coupled LED (M740F2, Thorlabs) coupled to a ferrule-terminated patch cable (200-μM core, 0.22 N.A., Thorlabs) attached to a fiber optic cannula (200-μM core, 0.22, Thorlabs). The cannula was glued to the ventral side of the holder and positioned approximately 135° from the front of the fly to be unobtrusive to the fly’s visual field. Throughout the experiment, saline bubbled with 95% O_2_ and 5% CO_2_ was superfused over the fly using a gravity fed pump at a rate of 2 mL/min. Whole-cell current-clamp recordings were performed using an Axopatch 200B amplifier with a CV-203BU headstage (Molecular Devices). Data from the amplifier were low-pass filtered using a 4-pole Bessel low-pass filter with a 5 kHz corner frequency, then acquired on a data acquisition card at 20 kHz (NiDAQ PCIe-6363, National Instruments). The liquid junction potential was corrected by subtracting 13 mV from recorded voltages^61^. Membrane potential data was then resampled to a rate of 1kHz for ease of use and compatibility with behavioral data. To estimate baseline membrane voltage (Fig. 5e,f,g) we removed spikes from voltage traces by median-filtered using a 50 ms window and lightly smoothed using the ‘smoothdata’ function in Matlab (loess method, 20 ms window). For all electrophysiology experiments in the mixed PFL2 & PFL3 line we would record from only one cell per fly. During recordings the cell would be filled using internal containing neurobiotin citrate so that we could collect the brain and visualize the cell morphology in order to determine its identity using the protocol described below in the Immunohistochemistry section.

### Spherical treadmill and locomotion measurement

Experiments used an air-cushioned spherical treadmill and machine vision system to track the intended movement of the animal at all times. The treadmill consisted of a 9 mm diameter ball machined from foam (FR-4615, General Plastics), sitting in a custom-designed concave hemispherical holder 3D printed from clear acrylic (Autotiv). The ball was floated with medical-grade breathing air (Med-Tech) through a tapered hole at the base of the holder using a flow meter (Cole Parmer). For machine vision tracking, the ball was painted with a high-contrast black pattern using a black acrylic pen, and illuminated with anIR LED (880 nm for 2-photon experiments; M880L3, Thorlabs, or 780 nm for electrophysiology experiments; M780L3, Thorlabs). Ball movement was captured online at 60 Hz using a CMOS camera (CM3-U3-13Y3M-CS for 2-photon imaging, or CM3-U3-13Y3C-CS for electrophysiology, Teledyne FLIR) fitted with a macro zoom lens InfiniStix 68-mm 0.66× for electrophysiology for 2-photon, InfiniStix 94-mm 0.5× for electrophysiology). This camera faced the ball from behind the fly (at 180°). Machine vision software (FicTrac v2.1) was used to track the position of the ball^49^ in real time. We used a custom Python script to output the forward axis ball displacement, yaw axis ball displacement, forward ball displacement, and gain-modified yaw ball displacement to an analogue output device (Phidget Analog 4-Output 1002_0B) and recorded these signals along with other experimental timeseries data on a data acquisition card at 20kHz (NiDAQ PCIe-6363) card at 20 kHz. The gain-modified yaw ball displacement voltage signal was also used to update the azimuthal position of the visual cues displayed by the visual panorama.

### Visual panorama and visual stimuli

To display visual stimuli, we used a circular panorama built from modular square (8×8 pixel) LED panels^50^. The circular arena was 12 panels in circumference and 2 panels tall. To accommodate the ball-tracking camera view and the light source the upper panel 180° behind the fly was removed. In all experiments, the modular panels contained blue LEDs with peak blue (470 nm) emission; blue LEDs were chosen to reduce overlap with the GCaMP emission spectrum. For calcium imaging experiments, four layers of gel filters were added in front of the LED arena (Rosco, R381) to further reduce overlap in spectra. For electrophysiology experiments, only 2 layers of gel filters were used. On top of the gel filters in both cases we added a final diffuser layer to prevent reflections (SXF-0600, Snow White Light Diffuser, Decorative Films). The visual stimulus displayed was a bright (positive contrast) 2-pixel-wide (7.5°) vertical bar. The bar’s height was the full 2-panel height of the area (except for −165 to +165° behind the fly with a single visual display panel, where the bar was half this height). The bar intensity was set at a luminance value of 4 with a background luminance of 0 (maximum value 15).

The azimuthal position of the bar was controlled during closed-loop experiments by the yaw motion of the ball (see Spherical treadmill and locomotion measurement). For all experiments, a 0.7x yaw gain was used, meaning that the visual cue displacement was 0.7× the ball’s yaw displacement. For calcium imaging and electrophysiology experiments the cue was instantaneously jumped every 60 seconds by ±90° or 180°. Immediately following the jumps, the cue would continue to move in closed loop with the fly’s movements. We recorded the position of the cue during experiments using analog output signals from the visual panels along with other experimental timeseries data on a data acquisition card at 20kHz (PCIe-6363, National Instruments). We converted analog signals from the visual panels into cue position in pixels during offline analysis. Cue positions were then converted into head direction as follows: 0° represents when the fly was directly facing the cue, 90° when the fly’s head direction was 90° clockwise to the cue, −90° when the fly was 90° counterclockwise, and 180° when the fly was facing directly away from the cue. These signals were lightly smoothed and values above 180 or below −180 were set to ±180°.

### Experimental trial structure

For all experiments prior to data collection, flies walked for a minimum of 15 minutes in closed loop with the visual cue. For calcium imaging experiments, data were collected in 10 minute trials. In each trial, the animal was in closed loop with a bright bar, and every minute the cue jumped to a new location relative to its current one, alternating between +90, 180, and −90, in order. In between trials during calcium imaging experiments, there was 30 seconds of darkness. During this inter-trial period in electrophysiology experiments, the flies were provided a light bar cue in closed loop. As these experiments were heavily dependent on spontaneously performed behavior, trials were run until the fly stopped walking or, in the case of electrophysiology experiments, the cell recording quality significantly decreased.

### Iontophoresis stimuli

Pipettes for iontophoresis were pulled from aluminosilicate capillary glass (OD 1.5 mm, ID 1.0 mm, Sutter Instrument Company) to a resistance of ∼75 MΩ using a horizontal pipette puller (P-97, Sutter Instrument Company). Pipettes were filled with a solution^62^ consisting of 10 mM ATP disodium in extracellular saline with 1 mM AlexFluor 555 hydrazine (Thermo Fisher Scientific) for visualization. This solution was stored in aliquots at −20C, thawed fresh daily, and kept on ice during the experiment. The tip of the iontophoresis pipette was positioned to be approximately in the medial region of the PB every trial. During experimental trials, we simultaneously recorded from a PFL2 neuron. During control trials, we recorded from unidentified neurons with somas in the same approximate region as PFL2 somas (medial area dorsal to the PB). Pulses of ATP were delivered using a commercial dual current generator iontophoresis system (Model 260, World Precision Instruments). Hold current was set to 10 nA to prevent solution leakage, and a current of −200nA was used for ejection. Visual confirmation of ATP ejection following current pulses was obtained before and after each trial. For the duration of the 10 minute trial period, flies were provided a light bar cue that moved in closed loop with their rotational movements, as described previously. Throughout the trial, pulses were delivered every 30 seconds with a length of 100, 200, 300 and 500 ms, repeating in that order.

### Immunohistochemistry

Brains were dissected from female flies 1-3 days post-eclosion in *Drosophila* external saline and fixed in 4% paraformaldehyde (Electron Microscopy Sciences, #15714) in phosphate-buffered saline (PBS, Thermo Fisher Scientific, 46-013-CM) for 15 min at room temperature. Brains were then washed with PBS before adding a blocking solution containing 5% normal goat serum (Sigma-Aldrich, #G9023) in PBST (PBS with 0.44% Triton-X, Sigma-Aldrich, #T8787) for 20 min. Brains were then incubated in primary antibody with blocking solution for ∼24 h at room temperature, washed in PBS, and then incubated in secondary antibody with blocking solution for ∼24 h at room temperature. Primary and secondary antibodies were protocol-specific (see below). Brains were then rinsed with PBS, and mounted in antifade mounting medium (Vectashield, Vector Laboratories, # H-1000) for imaging. For MCFO protocols, a tertiary incubation step for ∼24 h at room temperature and wash with PBS was performed prior to mounting. Mounted brains were imaged on a Leica SPE confocal microscope using a 40X 1.15 NA oil-immersion objective. Image stacks comprised 50 to 200 z-slices at a depth of 1 μm per slice. Image resolution was 1,024×1,024 pixels. For visualizing Gal4 expression patterns, the primary antibody solution contained chicken anti-GFP (1:1,000, Abcam, # ab13970) and mouse anti-Bruchpilot (1:30, Developmental Studies Hybridoma Bank, nc82). The secondary antibody solution contained Alexa Fluor 488 goat anti-chicken (1:250, Invitrogen, #A11039) and Alexa Fluor 633 goat anti-mouse (1:250, Invitrogen, #A21050). For visualizing cell fills after whole-cell patch clamp recordings, streptavidin::Alexa Fluor 568 (1:1000, Invitrogen, #S11226) was added to the primary and secondary solutions.

For MultiColor FlpOut (MCFO)^51^, the primary antibody solution contained mouse anti-Bruchpilot (1:30, Developmental Studies Hybridoma Bank, nc82), rat anti-Flag (1:200, Novus Biologicals, #NBP1-06712B), and rabbit anti-HA (1:300, Cell Signal Technologies, #NBP106712B). The secondary antibody solution contained Alexa Fluor 488 goat anti-rabbit (1:250, Invitrogen, #A11039), ATTO 647 goat anti-rat (1:400, Rockland, #612-156-120), and Alexa Fluor 405 goat anti-mouse (1:500, Invitrogen, #A31553). The tertiary antibody solution contained DyLight 550 mouse anti-V5 (1:500, Bio-Rad, #MCA1360D550GA).

### Processing calcium imaging data

The calcium imaging dataset comprised 23 flies expressing GCaMP under the control of the PFL3+2 split-Gal4 line, and 33 flies expressing GCaMP under the control of the PFL2 split-Gal4 line. Rigid motion correction in the x, y, and z axes was performed for each trial using the NoRMCorre algorithm^52^. Each region of interest (ROI) was defined across the z-stack. For each ROI ΔF/F was calculated with the baseline fluorescence (F) defined as the mean of the bottom 10% of fluorescence values in a given trial (600 s in length). From this measurement a modified z-score was calculated using the median absolute deviation (MAD) normalized difference from the median, which we refer to as the z-scored ΔF/F:

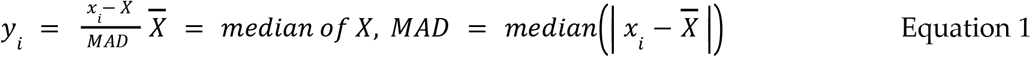

For PB imaging, 10 ROIs were defined, one for each of the 10 glomeruli occupied by PFL2 dendrites and defined to be approximately the same width and without overlap, constrained by estimated anatomical boundaries. For FB imaging, 9 ROIs were defined for PFL2 neurites corresponding to the 9 columns spanning the horizontal axis of the FB. ROIs were approximately the same width without overlap. For LAL imaging, two ROIs were defined, one for the left LAL and one for the right.

### Processing locomotion and visual arena data

Position of the spherical treadmill was computed online using machine vision software, and output as a voltage signal for acquisition. For post-hoc analysis, the voltage signal was converted into radians and unwrapped. Signals were then low-pass filtered using a second order butterworth filter with 0.003 corner frequency, and downsampled to half the ball-tracking update rate.Velocity was calculated using the MATLAB gradient function. Artifactually large velocity values (>20 rad/s) were set to 20 rad/s, and timeseries were then smoothed using the smooth function in MATLAB (using the loess method with an 33 ms window) and resampled to 60 Hz, the ball-tracking update rate. Forward and sideways velocities were then converted to mm/s while yaw velocity was converted to deg/s.

During calcium imaging, we acquired a signal from our imaging software indicating the end of each volumetric stack on the same acquisition card as on-line ball tracking signals. These imaging time points were then resampled to the ball-kinematic data update rate of 60 Hz and allowed us to align the acquired volumes. Electrophysiology data were collected on the same acquisition card as on-line ball tracking signals, so alignment was not required, however ball-tracking data were resampled to 1 kHz to match the sampling rate of the electrophysiology data.

### Computing inferred goal direction and walking straightness across trials

Head direction (θ) and head direction vector strength (*ϱ*) were calculated for every datapoint over each entire trial using a 30 second window centered on each datapoint index. Here we excluded datapoints where the fly’s cumulative speed (forward+sideways+yaw) was <1.5 mm/s. At values below this threshold the fly is essentially standing still, so including these timepoints might result in an overestimation of the fly’s internal drive to maintain its head direction. We also excluded timepoints within 5 seconds after a cue jump; this was to avoid underestimating the fly’s internal drive to maintain its head direction, as these points represent a forced deviation from the angle the flies were attempting to maintain. If no datapoints within the 30 second window satisfied these requirements they were excluded from further analyses. For included datapoints, head directions were treated as unit vectors. The average of these vectors was found to give the *ϱ* and θ values for each index. A *ϱ* value of 1 indicates the fly maintained the same head direction for the entire window, while a value of 0 indicates the fly uniformly sampled all possible head directions during the window. Fig. 1i shows mean *ϱ* and θ values from each trial.

### Categorizing behavior and inferring the fly’s internal goal

To categorize datapoints as part of straight or not-straight walking segments, we used a set threshold followed by a merge thresholded on segment duration (Fig. 4a). We empirically set the threshold for being defined as straight walking using a *ϱ* value (0.88 for imaging, 0.85 for electrophysiology). Datapoints continued to be categorized as straight walking until *ϱ* fell below that threshold. To form the “segments” used in further analysis, we merged any of the resulting straight walking epochs where intervening non-straight segments were less than 5 seconds.

We then calculated θ and *ϱ* for each of these segments, and used the mean θ value as the inferred goal head direction. For all analyses segments were discarded if *ϱ* was equal to 1, as this indicated the panels had not been initiated correctly and that the cue had remained in a single location for the duration of the trial. Segments were also discarded if the fly’s cumulative velocity was not above a threshold of 1.5 rad/s for at least 2 seconds. For population analyses shown in Fig. 1j (upper plot), 2e, and Fig. 3 all remaining segments were used regardless of *ϱ* or straightness category. For population analyses shown in Fig. 4b,c segments categorized as straight walking with a *ϱ* value above 0.5 were used to calculate the upper plot in both panels while segments categorized as not straight walking with *ϱ* values below 0.5 were used in the lower plots.

### Computing behavior around jump times

To analyze cue jumps shown in the lower plot of Fig. 1j, FIg. 4k, and in Extended Data Fig. 3 we calculated *ϱ* in the 10 seconds before and after each jump (+90, −90 or 180°), and measured the change in mean head direction. We categorized jumps as “corrected for” if the fly’s mean head direction returned to within 30° of its original location. We only included jumps where the fly had sufficient movement (cumulative velocity >1.5 mm/s for at least 3 seconds during the jump window) and was likely attempting to maintain a set head direction (*ϱ* was over 0.9 in the 10 seconds preceding the cue jump), and following the cue jump,. In Fig. 1j we combined jumps from PFL2/3 and PFL2 experiments prior to calculating mean and s.e.m. forward and yaw velocity. For the PFL2 and PFL2/3 specific plots in Fig. 4k and Extended Data Fig. 3 z-scored ΔF/F and bump amplitude were combined in the same manner, but only from the appropriate dataset.

### Computing average response to iontophoresis stimulation (2a,b)

For plots shown in Fig. 2a,b and Extended Data Fig. 5 data from the ±10 second period around each ATP pulse were averaged within individual flies to get the fly-averaged response to the 100 ms, 200 ms, 300 ms, and 500 ms pulses for the membrane potential, forward velocity, sideways velocity, and yaw velocity (each condition had at least 4 repetitions per fly). We then calculated the grand mean and s.e.m. across flies using these per-fly averages.

### Computing activity bump parameters (2c,d,f,g)

To track the amplitude and phase of PFL2 activity for analyses in Fig. 2c,d,e-g, Fig. 4f, and Extended Data Fig. 8 a sinusoid was fit independently to each timepoint of the z-scored ΔF/F activity across FB and PB imaging trials:

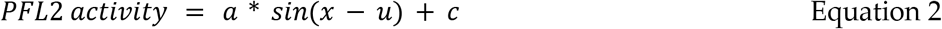

Here PFL2 activity is a vector of z-scored ΔF/F values at a single time point such that it has 10 bins if from a PB trial or 9, if from an FB trial corresponding to the number of ROIs specified for each region. Here, *ϱ*sets the phase of the sinusoid, *ϱ*is the vertical offset term, *ϱ*represents the bump amplitude and the position in brain space where the peak of the sinusoid is located defines the bump phase. A bump phase of +180° represents the rightmost position in the PB and FB while a phase of −180° represents the leftmost position.

### Computing change in bump phase vs change in head direction (2d)

We calculated the relative changes in PFL2 bump phase and head direction in 1.5 second bins shown in Fig. 2d and Extended Data Fig. 8. In each time window we took the difference between start and end points for HD or bump phase. Positive differences represent a clockwise shift while a negative difference represents a counterclockwise shift. The relationship between changes in HD and changes in bump phase was strongest when a 200 ms lag was implemented such that changes in bump phase lagged 200 ms behind changes in HD. The line of best fit for the relationship between the two variables was found with the polyfit and polyval functions. We then used the corrcoef function to find the correlation coefficients and p-value of the relationship. Indices where the adjusted r squared value of the sinusoidal fit for bump parameters were below 0.1 as well as those where fly was not moving were excluded from this analysis.

### Computing population activity as a function of behavior

To determine the relationship between neural activity and various behavioral parameters (Fig. 2e,f,g, Fig. 3, Fig. 4f-i, Extended Data Fig. 6) we binned conditioned data. Within each segment described above, indices with cumulative velocity < 1.5 mm/s were removed, and head directions were recalculated to be relative to the inferred goal head directions. Meaning that a negative value indicated that the fly was facing counterclockwise to its goal head direction, and a positive value indicated that the fly was facing clockwise to its goal head direction. The z-scored ΔF/F was then averaged within bins of 10 deg/s for yaw velocity, 1 mm/s for forward velocity, or 10° for head direction. For Fig. 3d,e, Fig. 4g,k and Extended Data Fig. 6, the sum or difference between right and left LAL activity was calculated per segment following binning. The mean and s.e.m. was then calculated across flies.

### Computing activity across brain space at different head directions

For the two example flies shown in Fig. 4i data were processed similarly to the above section, but in this case, we binned data from each ROI (PB glomeruli) individually across the entire non segmented trial to obtain the average response of each glomerulus across different head directions relative to the fly’s goal. The mean θ value associated with each index was calculated using a 60 second moving window centered on each index, as described previously (“Categorizing behavior into straight walking vs non-straight walking epochs & computing the inferred goal for each epoch.”). This θ value was taken as the inferred goal associated with that index. Head direction was divided into 4 bins centered on −90, 0, +90, and 180° from the goal with edges set ±45° from the centers. For each quadrant, the average binned z-scored ΔF/F value for each glomeruli was calculated with the rightmost glomeruli represented by angular position of +180 and the leftmost by −180.

### Computing goal amplitude as a function of head direction consistency

We used our computational model (see below, “Network model”, Equation 3) to predict amplitude of the goal signal across all FB and PB PFL2 trials using a nonlinear least squares fit (fit function in MATLAB), shown in Fig. 4j. To account for the arbitrary overall value of our fluorescence measurements, we added an overall activity offset term (*c*) to Equation 3. Head direction (θ) was resampled to match the sample rate for PFL2, and θ_g_ was calculated as previously described using a 30 second window centered on each index. Fit parameters were θ_0_, *A*, and *c*. Values of the offset (θ_0_) were bounded between 0 and 2*ϱ*, values of *A* had a lower bound of 0 but no upper bound, and *c* was unbounded. For every complete 10 minute trial, the vector strength was calculated for each index as described previously. Fitted values of *A* were binned by *ϱ* using 0.2 wide bins (lower limit of 0, upper limit 1) then averaged within each trial. We then averaged values of *A* across all trials for each fly, and calculated mean and s.e.m. across flies.

### Computing preferred head direction

To show preferred cell head direction in Fig. 5, we divided the estimated baseline membrane voltage (see **Patch-clamping**), into 20 degree bins based on the fly’s head direction. We considered the preferred head direction to be the value with the maximum binned membrane potential. The amplitude of the preferred head direction was calculated by taking the difference between the maximum and minimum binned membrane potential values.

### Analysis of IPSPs

To detect IPSPs for analyses in Fig. 5c and Extended Data Fig. 7, we focused only on jump trials where the fly was essentially immobile, to avoid any confounds associated with the membrane potential fluctuations in these cells that are associated with movement transitions (Extended Data Fig. 9). Action potentials were first removed from the voltage trace by median filtering the membrane potential with a 25 ms window, then lightly smoothing (smoothdata function in MATLAB, window size 20 ms, using the loess method). We then calculated the derivative of the membrane potential (gradient function in MATLAB), and found local minima corresponding to periods of rapid decreases in membrane potential (findpeaks function in MATLAB, peak distance of 20 ms, threshold determined for each cell). We also generated a detrended version of the membrane potential by subtracting the median filtered membrane potential (500 ms window), and found local minima (findpeaks function in MATLAB, peak distance of 20 ms, threshold determined for each cell). We categorized IPSPs as indices where a negative peak was detected from the derivative of the membrane potential trace within 30 ms before a negative peak in the baseline corrected trace.

### Computing change in IPSP parameters as a function of the change in head direction

To examine changes in IPSP parameters as a function of change in head direction, ±5 second windows around cue jumps in which the fly did not move for the entire 10 second period were used (Fig. 5a). All jumps fitting this category were analyzed for 6/7 neurons in this dataset, with data from each neuron plotted in a different color. The 7th neuron was not included as there were no cue jumps around which the fly was stopped for the entire 10 second window around the jump.

Detected IPSP frequency was calculated for the 5 seconds before or after the cue jump. The change in frequency before vs after jump was then compared to change in head direction relative to the cell’s preferred head direction produced by the cue jump. This was determined by first finding the absolute angular difference between head direction prior to the jump and the cell’s preferred head direction, computed as described previously, and doing the same for the new head direction following the cue jump. Then the pre cue jump value was subtracted from the post cue jump value. This means that a negative value indicated that the head direction got closer to the cell’s preferred head direction following the jump while a positive value indicated that the distance between the head direction and the cell’s preferred head direction increased following the jump. The change in IPSP frequency was then plotted against the change in the distance from the cell’s preferred head direction for each jump. MATLAB’s polyfit and polyval functions were used to find the line of best fit for the relationship between the two variables, while the corrcoef function was used to find the correlation coefficients of the relationship. Additionally we used an unbalanced two-factor ANOVA to determine the significance of the relationship between change in frequency and change in head direction.

### Computing change in preferred head direction amplitude between straight vs non straight segments

Each cell’s preferred head direction was calculated (see “Computing preferred head direction”) separately across all straight walking and all non straight walking segments (see above). The resulting preferred head direction curves are shown in Fig. 5e,f. Fig. 5g shows the amplitude of each cell’s preferred head direction (see above) across straight and non straight walking groupings. A two-sided Wilcoxon signed rank was used to determine the significance of the difference between the two distributions.

### Detecting and plotting neural activity around start/stop transitions

In Extended Data Fig. 9, movement transitions were defined as the time points at which the fly’s collective velocity passed a threshold value of 1.5 rad/s. These transitions were categorized as either start or stop transitions based on the following conditions. Start transitions were preceded by 2 seconds below the movement transition threshold and followed by 2 seconds above threshold (using a Gaussian-smoothed copy of the collective velocity). Stop transitions were followed by a 2 second period below threshold and preceded by a 2 seconds above threshold (using a gaussian-smoothed copy of the collective velocity). Start transitions were categorized as being near the cell’s preferred head direction if the head direction preceding the transition was within ±45° of the cell’s preferred head direction, and opposite to the preferred head direction if they were greater than ±135° away. Stop transitions were defined similarly but distance from preferred head direction was calculated using the head direction following the stop transitions. Membrane potential within a ±2 second window of these transitions were collected and smoothed using a moving average 75 ms in width to reduce the influence of more variable small fluctuations of membrane potential across the traces and averaged within each fly. The fly averaged traces are shown in the same color for each fly across the 4 categories.

### Neurotransmitter predictions

There are 25 PFL2 and PFL3 neurons in the hemibrain connectome, with over 100 pre-synapses associated with them. A recent effort to automatically infer transmitter identification from electron micrographs^23^ predicts that, of these, 12/12 PFL2 and 13/13 PFL3 neurons in the hemibrain connectome are cholinergic (Alexander S. Bates, personal communication). This neural network method predicts transmitters on a per-synapse basis, with an error rate that varies with cell and transmitter type. For PFL2 and PFL3 neurons, 7319 of 9870 (74%) confident presynapses (confidence score >= 0.5) were predicted as cholinergic. The second most commonly predicted transmitter was glutamate (11%). This work used 3094 hemibrain neurons in its ground truth data to train the model. These ground truth neurons could all be linked to cell types that had been identified as cholinergic using light microscopy pipelines and anti-body staining or RNA-seq. Among this ground-truth population, 73% of presynapses are correctly predicted as cholinergic. We, therefore, consider these prediction results to be strong evidence that PFL neurons use acetylcholine as their primary transmitter and are therefore excitatory

### Network model

Our model shares features with several other recent models^4,5,10,11,13^, However, it also incorporates information from the assignment of neurotransmitters based on machine classification^23^. Our model also differs from other models in a few important ways, as noted below.

To begin, head direction (θ) is encoded as a cosine-shaped spatial pattern of activity at the level of Δ7 cells in the protocerebral bridge. Specifically, the connectome predicts that there are two complete side-by-side spatial cycles of activity in Δ7 cells. Δ7 cells inherit head direction information from EPG cells as two von Mises-shaped bumps; there is evidence that they then reformat this signal into two cosine functions^5,24^, so that downstream neurons inherit cosine-shaped activity patterns^53,54^. Δ7 cells provide most of the head direction input to PFL2 and PFL3 cells^5^. The spatial phase of the Δ7 activity pattern should have an arbitrary offset relative to the fly’s head direction, with different values of θ_0_ in different individuals, and at different times in the same individual, because this is true of EPG cells^20^. We define θ_0_ as the angular position of the EPG bump at a head direction of 0°. The output of Δ7 cells is shifted by 180° versus the output of EPG cells, but Δ7 cells are also inhibitory; thus, we can model both EPG output and Δ7 output as cos(*x*-θ_0_+θ) over the spatial interval [0, 2*x*], where *x* is the horizontal distance from the left edge of the protocerebral bridge (over “neural space”). Note that the peak of activity moves rightward in the protocerebral bridge as head direction rotates leftward (counterclockwise)^55,56^. Negative values of this cosine function can be interpreted as values below some baseline or mean value.

PFL3 cells can be divided into two groups (PFL3>R and PFL3>L) based on whether they project to right or left descending neurons. The head direction maps in these two PFL3 groups are shifted ±67.5° relative to the map in Δ7 cells, as indicated by their anatomy^58^ (not ±90° as reported previously^5,10,11^). We model the head direction input to PFL3>R cells as cos(*x*+67.5°-θ_0_+θ), and we model the head direction input to PFL3>L cells as cos(*x*-67.5°-θ_0_+θ). Head direction input to PFL cells arises mainly from Δ7 cells^5^.

PFL2 cells sample one full head direction map from the middle section of the protocerebral bridge. Therefore, their head direction map is offset by 180°, relative to the map in Δ7 cells. Thus, we model the head direction input to PFL2 cells as cos(*x*+180°-θ_0_+θ).

We model the representation of the goal direction (θ_g_) as another sinusoidal pattern, which is reasonable, because the goal direction can be thought of as just a special head direction, and head direction is represented as a sinusoid. Because PFL2, PFL3>R, and PFL3>L cells receive almost identical inputs in the fan-shaped body, we assume the goal signal is the same in all these cell types. The phase of this sinusoid must be defined in terms of the system’s current representational coding frame. Thus, we model the goal input to PFL2 and PFL3 cells as *Aϱ*cos(*x*+θ_g_-θ_0_). Note that the peak of activity in goal cells will move leftward as the goal rotates rightward. Note also that, if there is a change in the offset of the head direction system (θ_0_), the goal representation must also be reformatted. The amplitude of goal input (*A*) can be interpreted as the strength of the goal’s influence over the organism’s behavior.

To obtain PFL activity levels, we sum their head direction inputs and their goal inputs, and we pass the result through an Exponential Linear Unit or ELU. This gives us three population activity patterns:

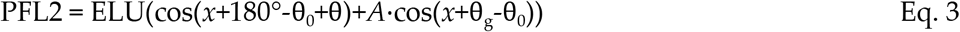

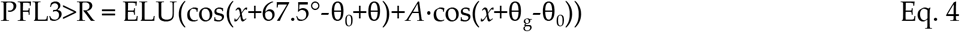

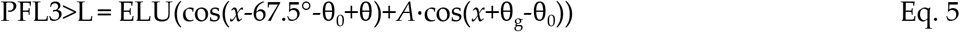

We must use a nonlinear function here, or else the goal will have no influence on the right-left difference in PFL3 activity (ΣPFL3>R - ΣPFL3>L). We choose to use an ELU unit because it is biologically highly plausible (as a “soft” rectifying nonlinearity like that observed in many cells^57^) and it is a good fit to our data. Note that other models have assumed a multiplicative^10^ or divisive nonlinearity^11^. Equations 3-5 were used to generate the model results in all figures except Fig. 3b-d.

To generate a neural-behavior feedback loop (in Fig. 3b-d), we model the effect of the PFL3 ensemble on the fly’s rotational velocity. We then take the fly’s rotational velocity at each time point and we feed it back into the brain’s head direction representation at the next time point. To model rotational velocity, we consider the fact that each PFL3 population synapses directly onto a DNa02 descending neuron^4,5^, and the right-left difference in DNa02 activity is known to be proportional to the fly’s rotational velocity, on average^4^. We obtained machine neurotransmitter classifications^23^ and found that PFL3 cells are cholinergic (confirmed for 13 of the 24 PFL3 cells in the hemibrain1.2.1^58^ dataset). This means that PFL3 cells are excitatory, not inhibitory as assumed elsewhere^5^. Thus we model the “direct” PFL3 contribution to rotational velocity as ΣPFL3>R - ΣPFL3>L. Note that positive values of rotational velocity denote rightward turns, and negative values denote leftward turns, in accordance with the convention in our field. Thus, in our model, PFL3>R cells promote rightward turning (not leftward turning, as assumed previously^5,13^). In Figure 1, we consider only this “direct” influence of PFL3 cells on steering.

At the same time, PFL3 cells also make equally strong connections onto DNa03, which is one of the strongest excitatory inputs to DNa02^5,28,29^. Thus, PFL3 cells are positioned to influence steering via a “direct pathway” and an “indirect pathway”. PFL2 cells do not connect directly to DNa02, but they make an anatomically massive connection onto DNa03, meaning they are positioned to scale the signals that flow through the indirect pathway. We found that PFL2 cells are also cholinergic and thus excitatory (again based on machine neurotransmitter classifications^23^, confirmed for 12 of the 12 PFL2 cells in the hemibrain1.2.1^58^ dataset). Thus, we model the influence of the indirect pathway as the total activity of PFL2 cells, multiplied by the right-left difference in PFL3 activity. In Figure 4, we therefore model the fly’s rotational velocity as the summed influence of both pathways:

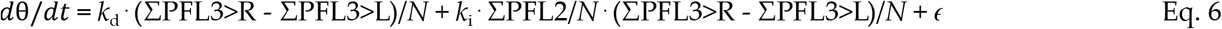

where *N* is the number of PFL neurons per population, *k*_d_ and *k*_i_ are constants and *ε* is a randomly fluctuating term that accounts for other sources of steering drive — e.g., sensory pathways that bypass the head direction system. We use *k*_d_ =100°/s and *k*_i_=2000°/s because these values generate approximately realistic steering behavior while avoiding oscillatory dynamics. We use *N*=360 to generate smooth curves over neural space, but it should be noted that the number of neurons in each PFL population is actually 12, according to hemibrain1.2.1^5^. We draw *ε* from a Gaussian distribution, then we low-pass filter *ε*(*t*) at 0.1 Hz, and we rescale *ε*(*t*) to enforce a standard deviation of 0.1°/s (Fig. 3b) or 100°/s (Fig. 3c,d).

## References

1. Schöne, H. Spatial Orientation: The Spatial Control of Behavior in Animals and Man. (Princeton University Press, 2014).

2. Knierim, J. J. & Zhang, K. Attractor dynamics of spatially correlated neural activity in the limbic system. Annu. Rev. Neurosci. 35, 267–285 (2012).

3. Hulse, B. K. & Jayaraman, V. Mechanisms underlying the neural computation of head direction. Annu. Rev. Neurosci. 43, 31–54 (2020).

4. Rayshubskiy, A. et al. Neural circuit mechanisms for steering control in walking Drosophila. bioRxiv 2020.04.04.024703; doi.org/10.1101/2020.04.04.024703, (2020).

5. Hulse, B. K. et al. A connectome of the Drosophila central complex reveals network motifs suitable for flexible navigation and context-dependent action selection. eLife 10, (2021).

6. Bennett, S. The development of servomechanisms. in A history of control engineering, 1800-1930 (Peregrinus, 1979).

7. Mittelstaedt, H. Control systems of orientation in insects. Annual Review of Entomology 7, 177–198 (1962).

8. Haferlach, T., Wessnitzer, J. & Mangan, M. Evolving a neural model of insect path integration. Adapt. Behav. (2007).

9. Stone, T. et al. An anatomically constrained model for path integration in the bee brain. Curr. Biol. 27, 3069–3085.e11 (2017).

10. Dan, C., Kappagantula, R., Hulse, B. K., Jayaraman, V. & Hermundstad, A. M. Flexible control of behavioral variability mediated by an internal representation of head direction. bioRxiv 2021.08.18.456004 (2022).

11. Matheson, A. M. M. et al. A neural circuit for wind-guided olfactory navigation. Nat. Commun. 13, 4613 (2022).

12. Le Moël, F., Stone, T., Lihoreau, M., Wystrach, A. & Webb, B. The Central Complex as a Potential Substrate for Vector Based Navigation. Front. Psychol. 10, 690 (2019).

13. Goulard, R., Buehlmann, C., Niven, J. E., Graham, P. & Webb, B. A unified mechanism for innate and learned visual landmark guidance in the insect central complex. PLoS Comput. Biol. 17, e1009383 (2021).

14. Sun, X., Yue, S. & Mangan, M. A decentralised neural model explaining optimal integration of navigational strategies in insects. eLife 9, (2020).

15. Touretzky, D. S., Redish, A. D. & Wan, H. S. Neural representation of space using sinusoidal arrays. Neural Comput. 5, 869–884 (1993).

16. Wittmann, T. & Schwegler, H. Path integration — a network model. Biol. Cybern. 73, 569–575 (1995).

17. Heinze, S. & Homberg, U. Maplike representation of celestial E-vector orientations in the brain of an insect. Science 315, 995–997 (2007).

18. Sakura, M., Lambrinos, D. & Labhart, T. Polarized skylight navigation in insects: model and electrophysiology of e-vector coding by neurons in the central complex. J. Neurophysiol. 99, 667–682 (2008).

19. Luigi Petrucco, Hagar Lavian, Vilim Stih, You Kure Wu, Fabian Svara, Ruben Portugues. A hindbrain ring attractor network that integrates heading direction in the larval zebrafish. in Computational and Systems Neuroscience Meeting T35. (2022).

20. Seelig, J. D. & Jayaraman, V. Neural dynamics for landmark orientation and angular path integration. Nature 521, 186–191 (2015).

21. Sun, X., Yue, S. & Mangan, M. How the insect central complex could coordinate multimodal navigation. Elife 10, (2021).

22. Lin, C.-Y. et al. A comprehensive wiring diagram of the protocerebral bridge for visual information processing in the Drosophila brain. Cell Rep. 3, 1739–1753 (2013).

23. Eckstein, N. et al. Neurotransmitter classification from electron microscopy images at synaptic sites in Drosophila. bioRxiv doi.org/10.1101/2020.06.12.148775 (2020).

24. Turner-Evans, D. B. et al. The neuroanatomical ultrastructure and function of a biological ring attractor. Neuron 108, 145–163 e10 (2020).

25. Green, J., Vijayan, V., Mussells Pires, P., Adachi, A. & Maimon, G. A neural heading estimate is compared with an internal goal to guide oriented navigation. Nat. Neurosci. 22, 1460–1468 (2019).

26. Giraldo, Y. M. et al. Sun navigation requires compass neurons in Drosophila. Curr. Biol. 28, 2845–2852.e4 (2018).

27. Warren, T. L., Weir, P. T. & Dickinson, M. H. Flying Drosophila melanogaster maintain arbitrary but stable headings relative to the angle of polarized light. J. Exp. Biol. 221, (2018).

28. Schretter, C. E. et al. Cell types and neuronal circuitry underlying female aggression in Drosophila. eLife 9, (2020).

29. Li, F. et al. The connectome of the adult Drosophila mushroom body provides insights into function. eLife vol. 9 Preprint at https://doi.org/10.7554/elife.62576 (2020).

30. Franconville, R., Beron, C. & Jayaraman, V. Building a functional connectome of the Drosophila central complex. eLife 7, (2018).

31. Mizumori, S. J. & Williams, J. D. Directionally selective mnemonic properties of neurons in the lateral dorsal nucleus of the thalamus of rats. J. Neurosci. 13, 4015–4028 (1993).

32. Driscoll, L. N., Pettit, N. L., Minderer, M., Chettih, S. N. & Harvey, C. D. Dynamic Reorganization of Neuronal Activity Patterns in Parietal Cortex. Cell 170, 986–999.e16 (2017).

33. Schoonover, C. E., Ohashi, S. N., Axel, R. & Fink, A. J. P. Representational drift in primary olfactory cortex. Nature 594, 541–546 (2021).

34. Marks, T. D. & Goard, M. J. Stimulus-dependent representational drift in primary visual cortex. Nat. Commun. 12, 5169 (2021).

35. Fisher, Y. E., Lu, J., D’Alessandro, I. & Wilson, R. I. Sensorimotor experience remaps visual input to a heading-direction network. Nature 576, 121–125 (2019).

36. Kim, S. S., Hermundstad, A. M., Romani, S., Abbott, L. F. & Jayaraman, V. Generation of stable heading representations in diverse visual scenes. Nature 576, 126–131 (2019).

37. Fisher, Y. E., Marquis, M., D’Alessandro, I. & Wilson, R. I. Dopamine promotes head direction plasticity during orienting movements. Nature in press (2022).

38. Ocko, S. A., Hardcastle, K., Giocomo, L. M. & Ganguli, S. Emergent elasticity in the neural code for space. Proc. Natl. Acad. Sci. U. S. A. 2018/11/30, E11798–E11806 (2018).

39. Heinze, S., Narendra, A. & Cheung, A. Principles of insect path integration. Curr. Biol. 28, R1043–R1058 (2018).

40. Collett, M. & Collett, T. S. How does the insect central complex use mushroom body output for steering? Curr. Biol. 28, R733–R734 (2018).

41. Webb, B. & Wystrach, A. Neural mechanisms of insect navigation. Curr Opin Insect Sci 15, 27–39 (2016).

42. Kamhi, J. F., Barron, A. B. & Narendra, A. Vertical lobes of the mushroom bodies are essential for view-based navigation in Australian Myrmecia ants. Curr. Biol. 30, 3432–3437.e3 (2020).

43. Buehlmann, C. et al. Mushroom bodies are required for learned visual navigation, but not for innate visual behavior, in ants. Curr. Biol. 30, 3438–3443.e2 (2020).

44. Grover, D. et al. Differential mechanisms underlie trace and delay conditioning in Drosophila. Nature 603, 302–308 (2022).

45. Kim, I. S. & Dickinson, M. H. Idiothetic path integration in the fruit fly Drosophila melanogaster. Curr. Biol. 27, 2227–2238 e3 (2017).

46. Rule, M. E. et al. Stable task information from an unstable neural population. eLife vol. 9 Preprint at https://doi.org/10.7554/elife.51121 (2020).

47. Cowan, N. J. et al. Feedback control as a framework for understanding tradeoffs in biology. Integr. Comp. Biol. 54, 223–237 (2014).

48. Wilson, R. I. & Laurent, G. Role of GABAergic inhibition in shaping odor-evoked spatiotemporal patterns in the Drosophila antennal lobe. J. Neurosci. 25, 9069–9079 (2005).

49. Moore, R. J. et al. FicTrac: a visual method for tracking spherical motion and generating fictive animal paths. J. Neurosci. Methods 225, 106–119 (2014).

50. Reiser, M. B. & Dickinson, M. H. A modular display system for insect behavioral neuroscience. J. Neurosci. Methods 167, 127–139 (2008).

51. Nern, A., Pfeiffer, B. D. & Rubin, G. M. Optimized tools for multicolor stochastic labeling reveal diverse stereotyped cell arrangements in the fly visual system. Proc. Natl. Acad. Sci. U. S. A. 112, E2967–76 (2015).

52. Pnevmatikakis, E. A. & Giovannucci, A. NoRMCorre: An online algorithm for piecewise rigid motion correction of calcium imaging data. J. Neurosci. Methods 291, 83–94 (2017).

53. Lu, J. et al. Transforming representations of movement from body-to world-centric space. Nature 601, 98–104 (2021).

54. Lyu, C., Abbott, L. F. & Maimon, G. Building an allocentric travelling direction signal via vector computation. Nature 601, 92–97 (2021).

55. Turner-Evans, D. et al. Angular velocity integration in a fly heading circuit. eLife 6, (2017).

56. Green, J. et al. A neural circuit architecture for angular integration in Drosophila. Nature 546, 101–106 (2017).

57. Anderson, J. S., Lampl, I., Gillespie, D. C. & Ferster, D. The contribution of noise to contrast invariance of orientation tuning in cat visual cortex. Science 290, 1968–1972 (2000).

58. Scheffer, L. K. et al. A connectome and analysis of the adult Drosophila central brain. eLife 9, (2020).

59. Tirian and Dickson, 2017, The VT GAL4, LexA, and split-GAL4 driver line collections for targeted expression in the Drosophila nervous system. http://dx.doi.org/10.1101/198648

60. Lima, S.Q., Miesenbock, G. (2005). Remote control of behavior through genetically targeted photostimulation of neurons. Cell 121(1): 141-152.

61. Gouwens, N. W. & Wilson, R.I. Signal propagation in Drosophila central neurons. J.Neurosci. 29, 6239-6249 (2009)

62. Liu WW, Wilson RI. Glutamate is an inhibitory neurotransmitter in the Drosophila olfactory system. Proc Natl Acad Sci U S A. 2013 Jun 18;110(25):10294-9. doi: 10.1073/pnas.1220560110. Epub 2013 May 31. PMID: 23729809; PMCID: PMC3690841.

